# Copy number variation profile-based genomic subtyping of premenstrual dysphoric disorder in Chinese

**DOI:** 10.1101/2021.02.08.430168

**Authors:** Hong Xue, Zhenggang Wu, Xi Long, Ata Ullah, Si Chen, Wai-Kin Mat, Peng Sun, Ming-Zhou Gao, Jie-Qiong Wang, Hai-Jun Wang, Xia Li, Wen-Jun Sun, Ming-Qi Qiao

**Author notes:** Correspondence: Hong Xue, Mingqi Qiao.

## Abstract

Premenstrual dysphoric disorder (PMDD) affects nearly 5% women of reproductive age. The symptomatic heterogeneity, along with largely unknown genetics, of PMDD have greatly hindered its effective treatment. In the present study, 127 Chinese PMDD patients of the ‘invasion’ and ‘depression’ subtypes clinically differentiated by us earlier were analyzed together with 108 non-PMDD controls for genome-wide copy number variations (CNVs). Germline genomic DNA samples from white blood cells were subjected to AluScan sequencing-based CNV profiling, which enabled clustering of patient samples readily into the V and D groups, dominated by the “invasion” and “depression” clinical subtypes, respectively; the CNVs obtained with 100-kb windows yielded two clusters that were correlated with these subtypes with a consistency of up to 89.8%. Diagnostic correlation- and frequency-based CNV features of either CNV-gain (CNVG) or CNV-loss (CNVL) that could differentiate between V and D subtypes were selected and analyzed. CNVG features located preferentially in S2-phase replicating regions and enriched with steroid hormone biosynthesis pathway of genes were found protective against PMDD. Moreover, machine learning employing the correlation-based CNV features could predict with >80% accuracy whether a genomic sample was D-type, V-type or control. In terms of their CNV profiles, the D- and V-types differed more from one another than from the controls, thereby providing a genomic basis for the clinical D-V subtyping of PMDD. Genome-wide profiling of CNVs, as a new approach to complex disease genetics, has revealed recurrent CNVs and genomic features beyond individual genes and mutations underlying PMDD clinical diversity.

## Introduction

Premenstrual dysphoric disorder (PMDD) is a syndrome that afflicts 5-10% of women in their reproductive years (1). The severity of the syndrome is typically highest just before the menstruation period, suggesting that the symptoms were linked to hormonal changes. This has been confirmed by the findings of premenstrual neurosteroid fluctuations, and alterations in the sensitivity of GABAA receptors to neurosteroids giving rise to mood instability (2, 3). Cortical gamma-aminobutyric acid (GABA) levels also declined during the menstrual cycle in healthy women but increased in women with PMDD from the follicular phase to the mid-luteal and late luteal stages (4). Furthermore, PMDD has been associated with the estrogen receptor alpha gene *ESR1* (5), and the ESC/E(Z) genes affecting the interactions of sex hormones with other genes (6). Five major contributors to the etiology of PMDD include: (1) genetic susceptibility; (2) progesterone and its metabolite ALLO; (3) estrogen, serotonin and brain-derived neurotrophic factor (BDNF); (4) brain structure and function; and (5) the hypothalamic-pituitary-adrenal axis and hypothalamic-pituitary-gonadal axis (7). The schizophrenia-associated SNPs in *GABRB2*, located in introns 8 and 9 near an AluYi6 insertion, have been associated with both schizophrenia and bipolar disorder (8, 9), heroin addiction (10), altruism (11), autism and mental retardation (12). Deletion of gabrb2 genes from knockout mice also brought about schizophrenic symptoms that were alleviated by the antipsychotic Risperidone (13). Recently, analysis of germline copy-number-variations (CNVs) at the nsv1177513 site in Exon 11, and the esv2730987 site in Intron 6, of *GABRB2* in PMDD and schizophrenia patients showed that CNV alterations at both esv2730987 and nsv1177513 were significantly associated with schizophrenia in Chinese and Germans as well as PMDD in Chinese (14). Moreover, subjects with different levels of susceptibility to cancer could be distinguished by means of diagnostic CNV marker features selected from the germline genomes with the application of machine learning (15).

It is recognized that the symptoms of PMDD are consistent with multiple clinical subtypes. A Delphi survey led to the proposal of three symptoms-based types of PMDD, *viz*. a predominantly physical type, a predominantly emotional type, and a mixed type (16); and DSM-V proposed that PMDD is defined by one or more of the symptoms of marked affective lability, marked irritability or anger, marked depressed mood and hopelessness, and marked anxiety and tension, plus at least one of seven other symptoms. At the School of Basic Medicine, Shandong University of Traditional Chinese Medicine, the medical records also pointed to at least two major types of PMDD, viz. an irritability-marked ‘invasion’ type (58.9%) and a depressive mood-marked ‘depression’ type (27.5%) (17). In view of the spectrum of PMDD symptoms, the objective of the present study was to enquire whether the two major clinical subtypes of PMDD could be corelated with genomic profiles. Through genome-wide CNV profiling by AluScan next-generation sequencing (18, 19), the results revealed two large clusters of CNV profiles that were highly correlated with the clinical “depression” and “invasion” subtypes. Furthermore, CNV-gain (CNVG) and CNV-loss (CNVL) features diagnostic of PMDD or each of the two clinical subtypes were uncovered among CNVs called from sequence windows of different sizes, which were variously distributed in genomic regions of different replication timing and overlapped with genes in various genetic pathways of potential clinical relevance. These results provided genomic verification for the invasion- and depression-subtypes employed by us previously (17), which corresponded to part of the complex symptoms stipulated by DSM-V (20) as diagnostic criteria for PMDD.

## Methods

### Clinical assessments

Clinical diagnosis of PMDD patients (P-type subjects) from asymptomatic controls (C-type subjects) was performed in accordance to the protocol in Diagnostic and Statistical Manual of Mental Disorders (DSM-IV) by two psychiatrists independently. The identifications of ‘depression-type’ and ‘invasion-type’ subjects were carried out as previously described (17).

### Genomic DNA samples

Peripheral white blood cell DNA samples were collected from PMDD patients and non-PMDD control subjects with approval by the institutional ethic committee of Shandong University of Chinese Medicine. The patients and healthy volunteers who participated in this study all signed the informed consent form. The samples consist of a control cohort of 108 subjects and a PMDD cohort of 127 cases. The latter cohort was further divided into the depression-subtype (71 cases) and invasion-subtype (56 cases). The subtypings of the 127 PMDD cases were given in Table S1.

### AluScan sequencing and CNV calling

Samples of ~0.1μg DNA were subjected to inter-Alu PCR amplification using the four Alu-consensual primers AluY278T18, AluY66H21, R12A/267 and L12A/8 (18). The 200 bp to ~6 kb amplicons in each sample were employed to build a library for sequencing on the Illumina platform with 100 bp paired-end reads. According to the standard framework, all the reads were mapped to reference human genome hg19 downloaded from UCSC by BWA, followed by base recalibration and local realignment by GATK (21). CNVs were called from the AluScan sequences with the method of AluScanCNV2 (19, 22) based on sequence windows of 50-500 kb in 50-kb increments on the 22 autosomes and the X chromosome. The CNV profiles of all 108 control and 127 PMDD subjects were available in Table S2.

### Clustering and grouping of patient samples based on CNV profiles

The profiles of CNVG and CNVL called from the 127 P-group samples using different CNV-calling window sizes were separately subjected to correlation analysis and hierarchical clustering with 1,000 bootstraps using the ‘pvclust’ R package (23). The derived correlation heatmaps as well as the CNVG-based and CNVL-based dendrograms obtained for each window size were employed to determine the two subgroups of CNV profiles using two different grouping methods for cross validation.

In the first method, *viz*. the straightforward *‘cutree’* method, the sub-clustering was carried out using the *‘culree’* function from the ‘dendextend’ R package (24) to cut each dendrogram into 2-8 sub-clusters (Figure S1). The DNA samples located in the sub-cluster populated with the highest number of clinical depression-type samples among all the sub-clusters was referred as D-type genomic samples; and the DNA samples located in the remaining sub-clusters were combined and referred as V-type genomic samples. In the second, or *‘semi-supervised* method, some branches on the dendrograms were first rotated around their respective nodes to bring the closely co-localized samples into tightly knit sub-clusters enclosed by black square boxes on the diagonal of each heatmap. Thereupon, all the samples within the same block box were all designated as D-type or V-type genomic samples depending on whether the majority clinical subtype of the samples were depression-type or invasion-type. The designated D- and V-type genomic samples derived using the two grouping methods for ten different window sizes are shown in Table S3 and exemplified by the blue and red branches in Figure 1 and Figure S2 respectively for the 100-kb CNV profiles.

**Figure 1.**
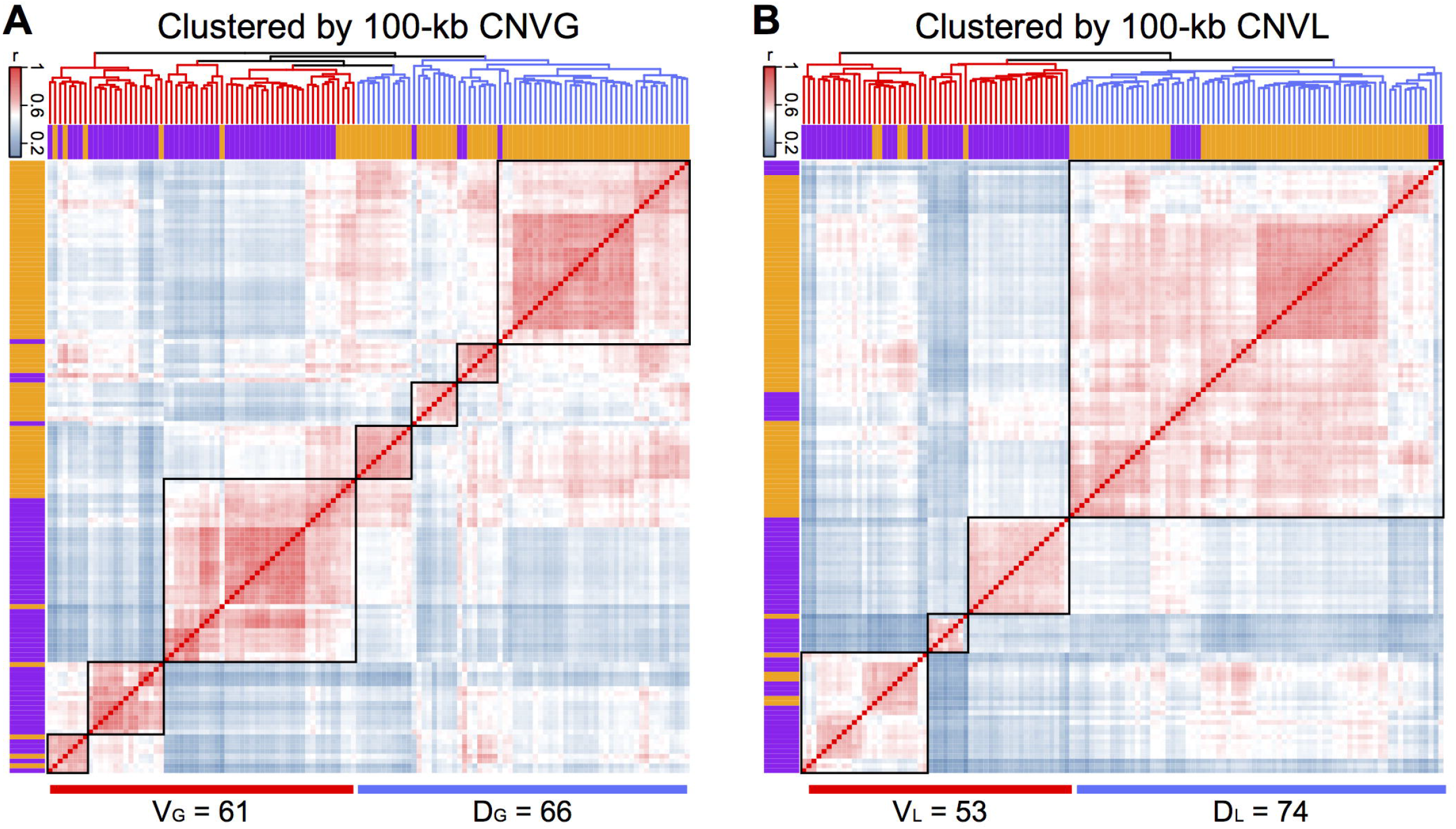
Hierarchical clustering of PMDD samples based on their pairwise similarities in genome wide CNV profiles. For all 127 P-group samples, all CNVs were identified from AluScan sequencing data with 100-kb non-overlapping scanning windows across the genome and used in the plots of similarity scores for CNVGs (A) and CNVL (B), respectively. The dendrograms on top of the heat maps were bootstrapped 1,000 times. The color of each square in the heat map indicates the correlation coefficient (r) of a pair of samples according to the blue-red thermal scale. The semi-supervised classification of samples based on (A) CNVGs and (B) CNVLs was indicated by the red dendrogram branches for V-type and blue ones for D-type genomes. The bands below the dendrograms and on the left-hand side of the heat maps portrayed the subtyping of PMDD samples based on clinical symptoms, with purple bands representing the clinically determined invasion subtype (n = 56) and orange bands the depression subtype (n = 71). Each of the square diagonal boxes in panels (A) and (B) enclosed a group of genomes with close correlations between each other in the group, such that they could be identified as a coherent block of genomes belonging to either the V-type or D-type CNV profiles depending on their enrichment in the invasion- or depression-subtype samples (see ‘Clustering of patient samples based on CNV profiles’ in Methods). Comparable heat maps obtained using sequence window sizes of 50 to 500 kb for CNV-calling are shown in Figure S3.

For either the *‘cutree”* method or the ‘*semi-supervised* method, let the number of CNV-based D-type samples that also belonged to the clinical depression-subtype be represented by *True_D_*, and the number of CNV-based V-type samples that also belonged to the clinical invasion-subtypes be represented by *True_V_*. Accordingly, the consistency (*Y*) between CNV-based classification and the clinical classification of PMDD patient samples could be estimated by:

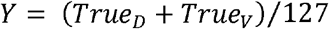

On this basis, the levels of consistency between CNV-based and clinical subtypings for the different CNVs called using different window sizes for both the *‘cutree’* and ‘*semi-supervised* methods are shown in Table S4.

### Selection of diagnostic CNV features

The selection of diagnostic CNV features was performed using either (a) correlation-based method or (b) frequency-based method as described (15). CfsSubsetEval from the Weka package was employed together with BestFirst search method to select the correlation-based diagnostic CNV features. Fisher’s exact tests were employed to select the frequency-based CNV features that showed significantly different occurrence frequencies between a pair of sample groups (e.g. P-vs-C or D-vs-V) with a false discovery rate (FDR) less than 0.01.

### Predictive subtyping of genomic samples by machine learning

Earlier, diagnostic germline CN-gains and CN-losses from leucocyte DNA samples of subjects with or without past episodes of cancers in tissues other than leucocytes were found to provide a useful basis to predict the propensity of the subject to cancer (15). Since the 127 P-group and 108 C-group DNA samples from PMDD and control subjects consisted of a mixture of D-type, V-type and C-type DNAs, the question arose whether it was possible to predict the typing of DNA samples between the D-vs-V, D-vs-C and V-vs-C choices employing the diagnostic CNVG and CNVL features obtained with the correlation-based method.

For example, in a choice between the P-vs-C types, a mixture of P- and C-type samples were randomly separated into a labeled Learning Band and an unlabeled Test Band, with equal or near equal number of samples in the two bands. Diagnostic CNVG and CNVL features were selected from the labeled Combined Learning Band with machine learning using the correlation-based method and employed to estimate the risk factor *R* for each DNA sample in the Test Band according to Eqn. 1.

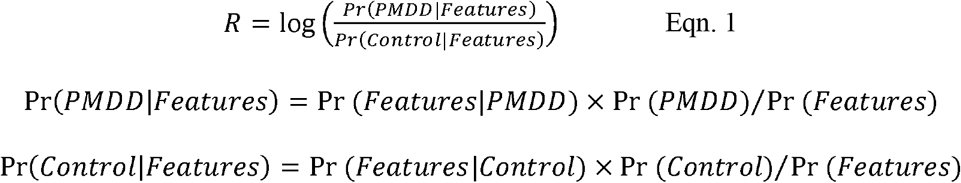

where Pr(PMDD|Features) was the posterior probability of membership in the PMDD group given the CNV data of a particular Test Band sample; Pr(Control|Features) was its posterior probability of membership in the Control group given the same test CNV data; Pr(Features|PMDD) was the likelihood function of the test CNV data given membership in the PMDD group; Pr(Features| Control) was the likelihood function of the test CNV data given membership in the Control group; Pr(PMDD) and Pr(Control) were the prior distributions of PMDD and Control samples respectively within the Learning Band; and Pr(Features) was the prior distribution of CNV-features among all the CNVs within the Learning Band.

For every sample in the Test Band, its value of R estimated using Eqn. 1 would predict whether the sample belonged to the Control group or PMDD group: it would predictively belong to Control group (*viz*. ‘non-PMDD’) if *R* < 0; belong to PMDD group if *R* > 0; or no prediction could be made if *R* = 0. For every PMDD sample in the Test Band, *R* >0 represented a ‘true’ prediction whereas *R* < 0 represented a ‘not true’ prediction. On the other hand, for any Control sample in the Test Band, *R* > 0 represented a ‘not true’ prediction whereas *R* < 0 represented a ‘true’ prediction. Accuracy of prediction was therefore given by:

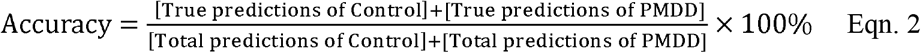

Repetition of this procedure 1,000 times would yield 1,000 Accuracy estimates, and in turn the Average Accuracy regarding the P-vs-C typing.

### Functional annotation of genes overlapping with diagnostic CNV features

By comparing the genomic coordinates of all the frequency-based diagnostic CNV features to those of the known genes retrieved from the R package ‘TxDb.Hsapiens.UCSC.hg19.knownGene’ version 3.2.2 (25), and considering any gene to be ‘overlapping’ with a CNV feature if any proportion of its sequence (from > 0% to 100% in 10% increments) coincided with part or all of the CNV feature, the list of CNV-overlapping genes obtained was uploaded to DAVID Bioinformatics Resources as a *test-list* employing the ‘RDAVIDWebService’ R package (26). All the known genes on chromosomes 1-22 and X were also uploaded as the *background-list*. Comparison of the two lists using the ‘getFunctionalAnnotationChart’ of the ‘RDAVIDWebService’ R package revealed gene pathways or categories, as defined in the GO, KEGG and INTERPRO databases, that were enriched with the *test-list* of genes among the *background-list* of genes. The pathways or categories yielding <0.05 Benjamini-corrected *p*-values were regarded to be significantly enriched in the genes on the *test-list* (Table S5).

### Genomic-feature content of diagnostic CNV features in different replication phases

DNA sequences on 22 autosomes and chromosome X were subject to replication-time segmentation according to Long & Xue (27). Briefly speaking, experiment-assessed replication timing of all 1-kb sequence windows in the genomes of fifteen human cell lines were retrieved from the ‘UW Repli-seq track’ in the UCSC Table Browser (28), and the representative replication phase of each sequence window was identified as one of the six types of sequencing segments (viz. G1b, S1, S2, S3, S4 and G2) based on their experiment-assessed replication timing in all fifteen human cell lines.

The density or intensity of genomic features were quantified as described in Ng et al (29) in the diagnostic CNV features and the non-diagnostic-CNV regions in each type of replication phase. The genomic feature content of diagnostic CNV features is indicated by the fold change of the density or intensity of the genomic feature in diagnostic CNV features relative to the non-diagnostic-CNV regions.

### Statistical analysis

All comparisons of CNV frequencies were conducted using Fisher’s exact tests, and the *p-* values were adjusted by false discovery rate for multiple comparisons. In functional annotation of genes, *p*-values from DAVID web service were subject to Benjamini-correction for multiple comparisons. When annotating the genes that overlapped with any diagnostic CNV feature, empirical *p*-values were estimated using Monte Carlo methods with 1,000 simulations to validate the significant gene pathways/categories based on the 50-, 100- or 450-kb size groups of CNV features. In each round of simulation, sequence windows of the same size as the targeted group of CNV features were randomly selected from chromosomes 1-22 and X, with the number of selected windows being equal to the average number of CNV features in the different type-comparisons to be analysed (see Table S6). For each simulation, the genes that overlapped with any of the selected sequence windows were functionally annotated. The empirical *p*-value of a targeted pathway was given by (r+1)/(n+1), where *n* = 1,000 and *r* = number of simulations that displayed significant enrichment (<0.05 Benjamini-corrected *p*-values) in the targeted pathway.

### Software for data processing and visualization

Data processing tasks were carried out using custom R codes, except that tasks requiring machine learning were processed using Weka package. All figures were drawn under R environment using the ‘ggplot2’ (30), ‘pheatmap’ (31) and ‘quantsmooth’ (32) packages, except for Figure 3 which was drawn using http://bioinformatics.psb.ugent.be/webtools/Venn/, and Figure 5 using Integrative Genomics Viewer 2.3.69 (33).

## Results

### Correlations between clinical diagnosis and CNV profiles

In order to examine whether there might be significant correlation between the clinical symptoms of PMDD patients and their germline CNV profiles, the CNVGs and CNVLs called from different sizes of sequence windows on the 127 P-type DNA samples, were subjected to hierarchical clustering in each instance. The CNVGs and CNVLs called from 100-kb sequence windows of the 71 depression-subtype and 56 invasion-subtype patient samples were segregated using the cutree and semi-supervised methods into distinct D-type and V-type clusters in the dendrograms as shown in Figure S2 and Figure 1 respectively. The clusters obtained from the CNVG dendrograms were designated as D_G_ and V_G_ clusters, and the clusters obtained from the CNVL dendrograms were designated as D_L_ and V_L_ clusters. Notably, the cutree method yielded 72 V_G_-type and 55 D_G_-type CNVG profiles with 81.10% consistency between the invasion-vs-depression clinical classification and the V-vs-D CNVG-based classification (Figure S2A); whereas the semi-supervised method yielded 61 V_G_-type and 66 D_G_-type CNVG profiles with 89.76% consistency between the invasion-vs-depression clinical classification and the V-vs-D CNVG-based classification (Figure 1A). Therefore, using either the cutree method or the semi-supervised method, the CNVG-based classification was highly correlated with the clinical symptom-based classification of the

PMDD genomes; this was likewise the case with the D_L_-type and V_L_-type CNVLs. Altogether, for the CNVGs and CNVLs in, the 50-500 kb window sizes, the cutree method yielded consistencies of 68-91%, and the semi-supervised method yielded consistencies of 88-98%, between the CNV-based and symptom-based classifications. The semi-supervised classifications of P-type samples based on CNVs called from 50-500 kb window sizes were available in Figure S3. These results demonstrated that both the CNVGs and CNVLs contributed to the etiology of the depression-type and the invasion-type symptoms. Moreover, the comparable results obtained using the cutree and semi-supervised methods confirmed the robustness of the CNV-symptom correlations. When the CNVGs or CNVLs called from 100-kb sequence windows of the 108 C-type control samples were subject to hierarchical clustering along with the P-type samples, ~40% of the C-type CNV profiles formed a tight sub-cluster and ~60% were dispersely distributed in the dendrogram, forming sub-clusters with the depression-subtype or invasion-subtype PMDD samples (Figure S4).

### Use of diagnostic CNV-features for predictive subtyping

The correlation between germline CNV profiles and clinical subtypes of PMDD suggests that it would be practicable to predict from the germline CNVs of women their propensity to develop PMDD, as well as the likely subtype of the PMDD clinical condition. Toward this objective, the method developed earlier by us through the use of diagnostic CNV-features selected with machine learning to assess a subject’s propensity for cancer (15) could be employed as described in ‘Selection of diagnostic CNV features’ under Method. Figure 2 shows the diagnostic CNV features selected by either the correlation method or the frequency method for prediction the propensity of a test subject’s germline CNVs for which of the P, C, V_G_, D_G_, V_L_ and D_L_ genomic groups: the P-type and C-type outcomes would be assessed based on PMDD symptoms; V_G_ and D_G_ would be based on the distinction between the V and D clusters in the CNVG dendrogram in Figure 1A; and V_L_ and D_L_ would be based on the distinction between the V and D clusters in the CNVL dendrogram in Figure 1B. The diagnostic CNVG and CNVL features selected using the correlation and frequency methods are given in Table S7.

**Figure 2.**
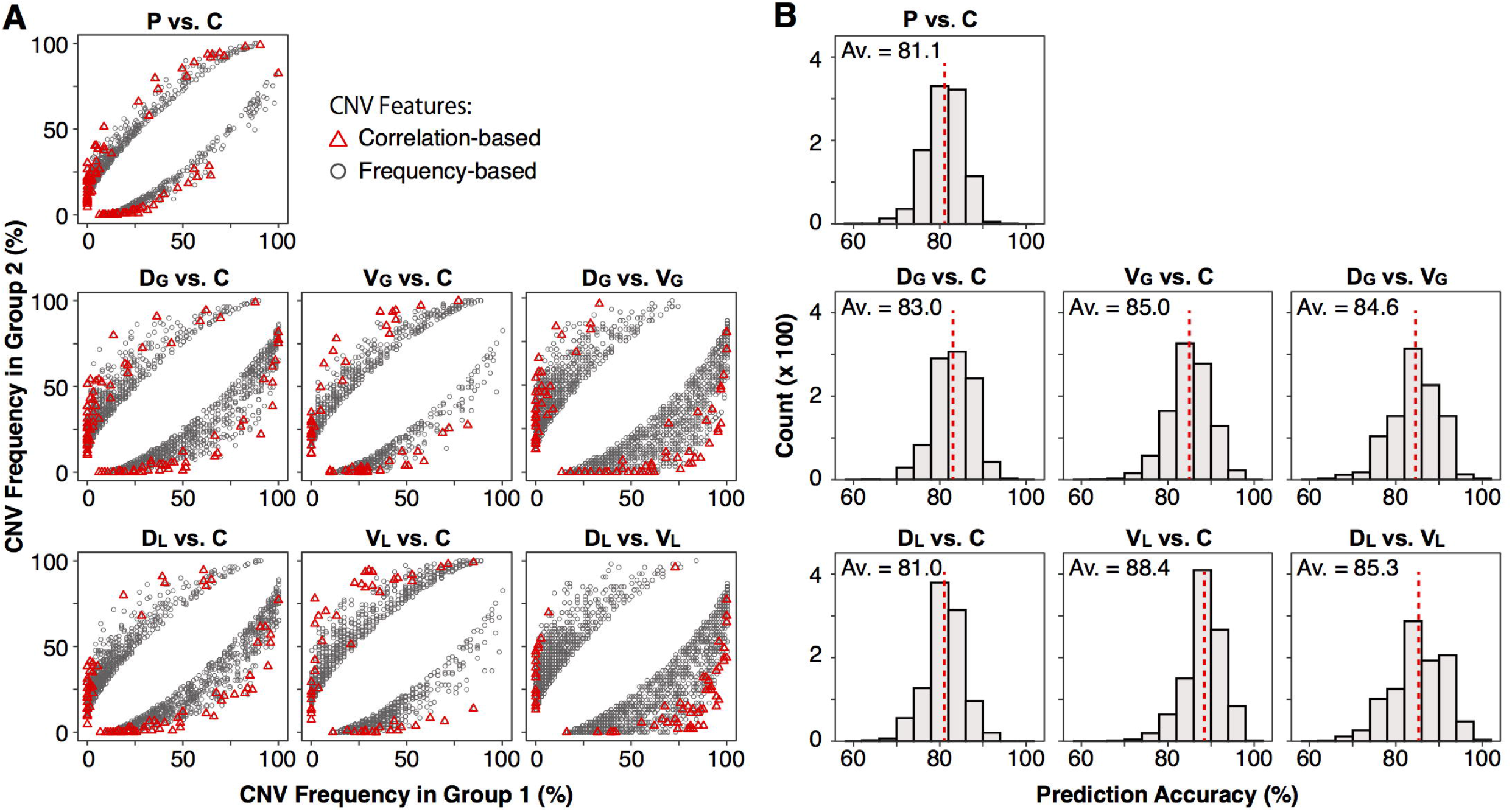
Occurrence frequencies of diagnostic CNV features and their prediction accuracies for seven pairs of sample groups. Panel (A) shows the frequency distribution of diagnostic CNV features for different pairs of sample groups. The x-axis represents the frequency of CNVs in the first-named group (Group 1 as shown on x-axis), and y-axis the frequency of CNVs in the second-named group (Group 2 as shown on y-axis) in a given pair of sample groups. Diagnostic CNV features with higher frequencies in Group 1 relative to Group 2 (located in lower right crescent) are referred to as ‘Group 1-favoring’ features, whereas diagnostic CNV features with higher frequencies in Group 2 relative to Group 1 (located in upper left crescent) are ‘Group 2-favoring’ features. Black circles are CNV features selected using the frequency-based method with FDR < 0.01 (Fisher’s exact tests), and red triangles are CNV features selected using the correlation-based method. Panel (B) shows the prediction accuracies (estimated using Eqn.2 in Methods) of sample classification in seven sample-pairs based on CNV features selected using the correlation method. For each of the seven pairs, prediction accuracy was estimated 1,000 times and the average accuracy (Av.) was given in the pertinent panel. Subscript G denotes that the D- or V-type samples were derived from the dendrogram of CNVGs (Figure 1A), while subscript L denotes that the D- or V-type samples were derived from the dendrogram of CNVLs (Figure 1B).

**Figure 3.**
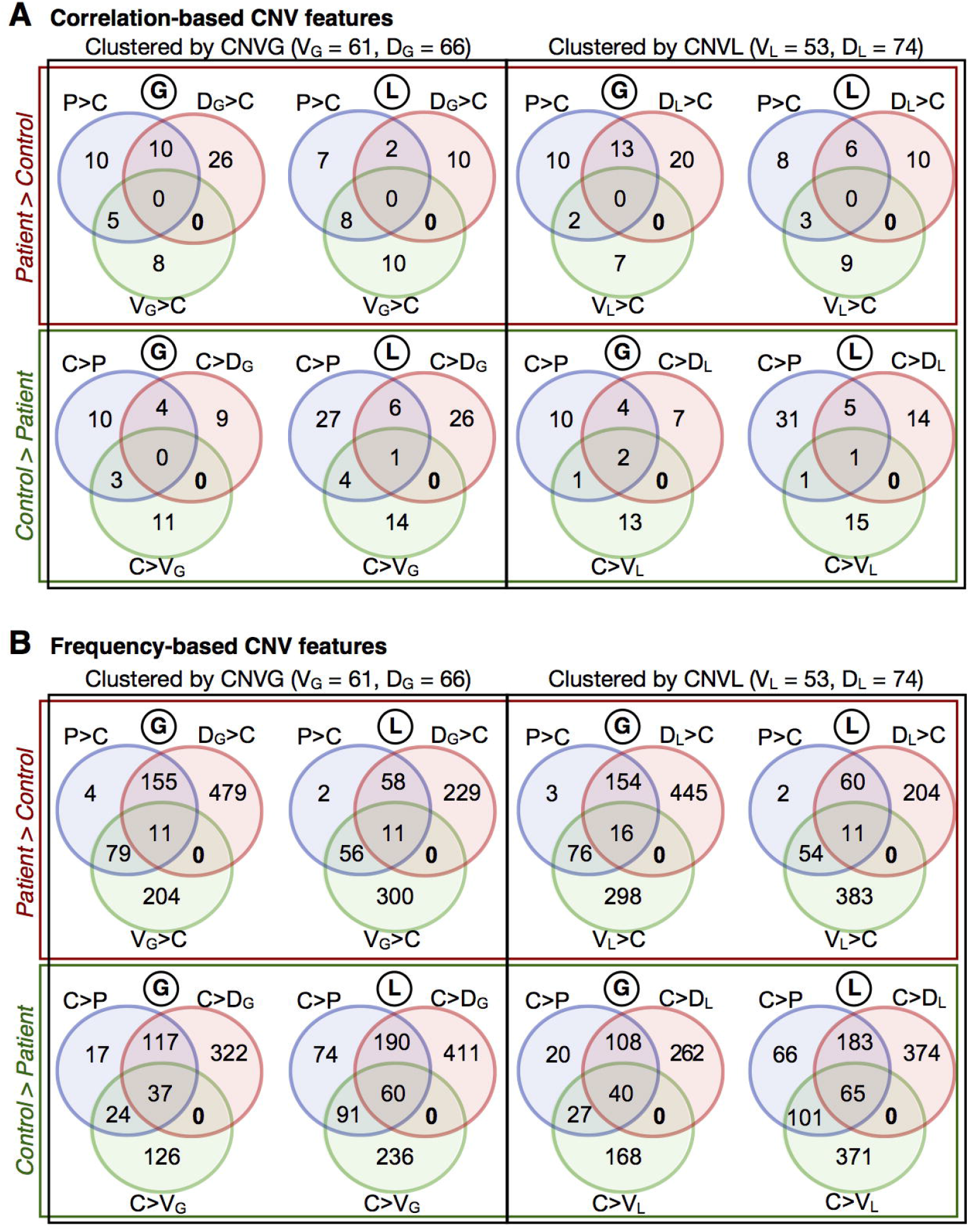
Overlaps between the diagnostic CNV features differentiating the two subtypes of PMDD collectively and individually from the control. CNV features identified using (A) correlation-based method, and (B) frequency-based method. Circled ‘G’ indicates CNVG features and circled ‘L’ indicates CNVL features. The ‘>’ and ‘<’ signs portray the relative frequencies of the CNV features for a pair of sample groups, e.g. P>C represents diagnostic CNV features that occurred in higher frequencies in P-group compared to C-group. Subscript G denotes that the D- or V-type samples were derived from the CNVG dendrogram in Figure 1A, whereas subscript L denotes that the D- or V-type samples were derived from the CNVL dendrogram in Figure 1B.

Figure 2A shows the sets of diagnostic CNV features selected using the correlation-based (red triangles) or frequency-based (black circles) method to enable a choice between a pair of genomic groups. For example, the D_G_-vs-C panel of Figure 2A contained a mixture of 66 D_G-_ type samples and 108 C-type samples. The diagnostic CNV features selected from the total of 144 samples by means of either the correlation method (red triangles) or the frequency method (grey circles) were distributed in a crescent near the y-axis and another crescent near the x-axis. Accordingly, any DNA sample in the mixture that was enriched with near-y diagnostic CNV features would be predicted to be endowed with a greater propensity for C-type over D_G_-type, whereas any DNA sample that was enriched with near-x diagnostic CNV features would be predicted to be endowed with a propensity for D_G_-type over C-type. In the D_G_-vs-C panel of Figure 2B, diagnostic CNV features selected using the correlated method was employed to predict the D_G_-vs-C nature in the 174-sample mixture as described under the ‘Predictive subtyping of genomic samples by machine learning’ section in Methods. After 1,000 trial runs, each with a random partition of the samples into an 87-sample Learning Band and an 87-sample Test Band, the average prediction accuracy obtained was 83.0%. Altogether, the seven panels in Figure 2B yielded average prediction accuracies ranging from 81.0% to 88.4%. Interestingly, the list of correlation-based CNV features useful for differentiating between the propensities toward the D and V subtypes (Table S8) showed that the CNV features biased in favor of V-type samples were mostly CNVL features (27/42 for V_G_ and 14/17 for V_L_). The accuracies of sample-classification predictions derived from the cutree method are available in Figure S5.

Favorable diagnostic CNV-features were often shared by more than one PMDD types, as indicated by the overlaps between the colored circles for the P-vs-C (blue), D-vs-C (red) and V-vs-C (green) comparisons in the Venn diagrams (Figure 3A and B). A range of CNV features were shared by all three kinds of circles, suggesting that they represented key CNV features differentiating between the control and PMDD patient samples (Table S9). Notably also, in all the panels in Figure 3, there was no CNV feature was shared only by the red circles for D-vs-C and the green circles for V-vs-C, which suggests that the CNV-features favoring the D-type genomes differed diametrically from the CNV-features favoring the V-type genomes. As well, there were more D-favoring CNVG features than CNVL features, but more V-favoring CNVL features than CNVG features.

### Genome-wide distribution of diagnostic CNV features

In order to have a global view of CNV profiles, the locations and replication timing of all frequency-based diagnostic CNV features, whether overlapping with any known genes or not, were plotted on Figure S6. The results showed that the CNV features were widely spread on all the somatic chromosomes and chromosome X. Chromosomes 4, 13, 18 21 and X were particularly abundant in CNV features that replicated in the G2 phase. Given the correlation between the clinical symptom-based typing of PMDD cases and the clustering of germline diagnostic CNV features, these CNV features could be useful guides in a search for some of genomic sites underlying PMDD.

In Figure 4, the distributions of the CNV features among DNA regions replicating at different cell cycle phases exhibited a number of characteristics: (a) In terms of the number of CNV features that differed between a pair of CNV-types, the P-vs-C panel (viz. P>C or P<C) gave rise to the smallest difference, whereas the D-vs-V pair (D_G_>V_G_ or D_G_<V_G_) gave rise to the largest difference; (b) the ratio of CNVL features relative to CNVG features (viz. L/G on chart) that favored the C-type over P-type were 1.32 for 50-kb CNV features, 2.13 for 100-kb ones and 2.05 for 450-kb ones, all greater than unity (Figure 4A); (c) the P-vs-C comparisons were suggestive of protective effects of smaller size CNVLs in the early replication phases and larger CNVLs in the later phases (Figure 4A); (d) the CNVLs captured by 50-kb windows included significantly more V-favoring than either C-favoring or D-favoring ones (L/G = 2.41 in Figure 4C and 1.80 in Figure 4D); (e) the CNVGs were significantly enriched in D-favoring features compared to C-favoring or V-favoring ones, whereas CNVLs were significantly enriched in V-favoring features compared to C-favoring or D-favoring ones; (f) D-vs-V comparisons suggest that V-type PMDD was correlated with smaller CNVG features belonging to the early replication phases and large CNVGs belonging to the later phases; (g) Large CNVG features were enriched in the G2-phase replicating sequences, especially among the features selected for the D-vs-C and D-vs-V comparisons (see G2-phase columns marked with red asterisks in Figure 4B and D); (h) More than half of the large G2-phase CNVG features in the C>D_G_ group are identical to those of the V_G_>D_G_ group, suggesting the shared genetic variations in G2 phase underlying V and C types; (i) Large CNVL feature were enriched in S3-phase replicating sequences in the C>V_G_ and D_G_>V_G_ groups but not in the V_G_>C or V_G_>D_G_ groups. The replication-phase distributions of CNV features obtained based on the D_L_- or V_L_-type samples derived from the CNVL dendrogram in Figure 1B were available in Figure S7.

**Figure 4.**
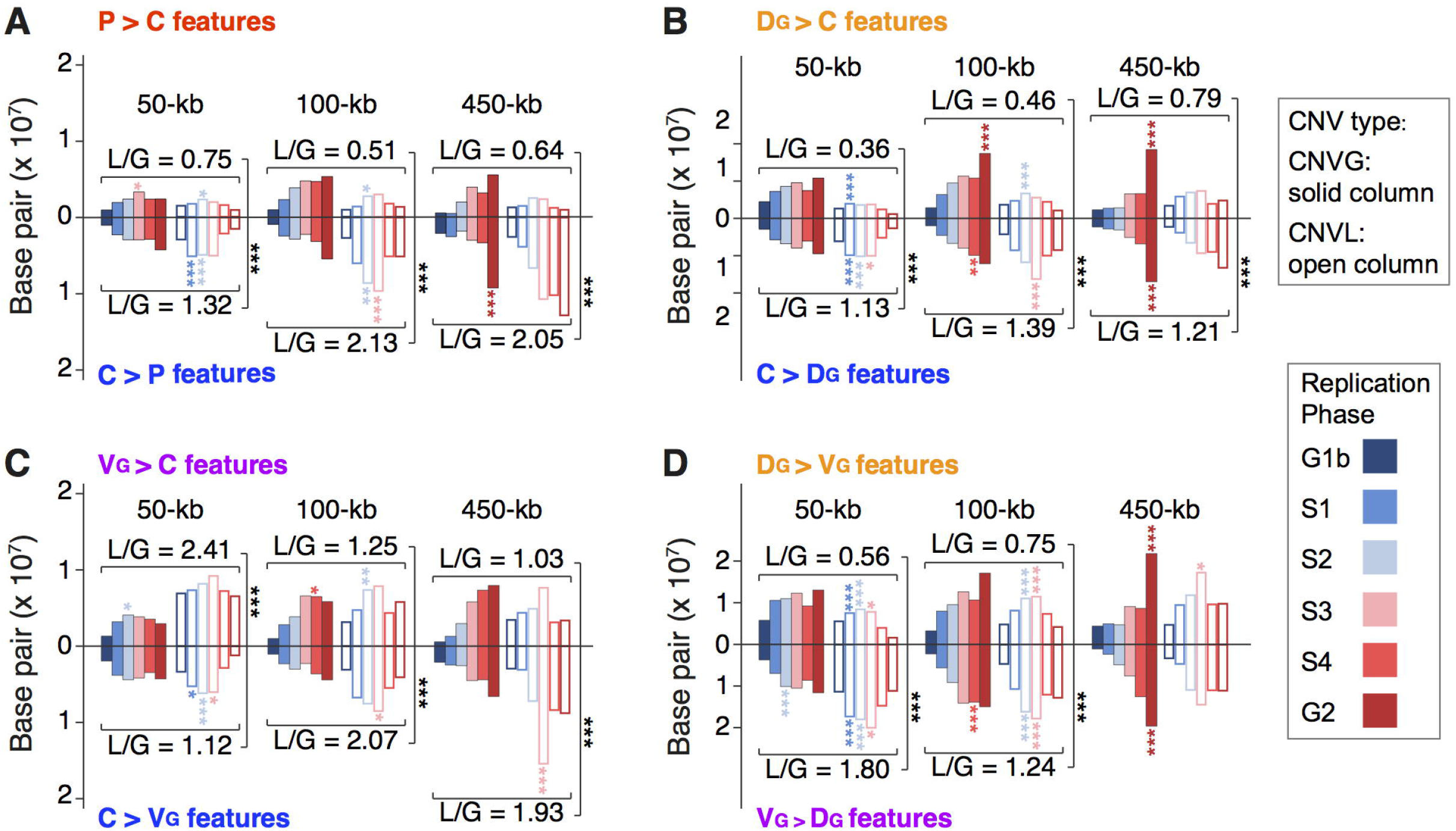
Distribution of frequency-based diagnostic CNV features among genomic sequences of different DNA replication phases. Number of base pairs of the CNV features called using 50, 100 and 450-kb windows for (A) P-vs-C, (B) D_G_-vs-C, (C) V_G_-vs-C, and (D) D_G_-vs-V_G_ groups. The solid bars represent CNVG features and hollow bars represent CNVL features in each panel. The replication phases G1b to G2 are color coded as shown. The ‘>’ or ‘<’ sign portrays larger or smaller frequencies of the CNV features in favor of the first-named group over the second-named one. L/G represents the ratio of the number of CNVLs over the number of CNVGs. Significant enrichment of CNV features in a particular replication phase in the genome is indicated by asterisks that are color coded according to the replication phase, or in black asterisks for comparison between an L/G value in the upper half of a panel and an L/G value in the lower half (Bonferroni-corrected, *** *p* < 0.005, ** *p* < 0.01, * *p* < 0.05). Numerical *p*-values are shown in Table S14. Subscript G denotes that the D- or V-type samples were derived from the CNVG dendrogram in Figure 1A. See Figure S7 for the results obtained based on the D- or V-type samples derived from the CNVL dendrogram in Figure 1B.

**Figure 5.**
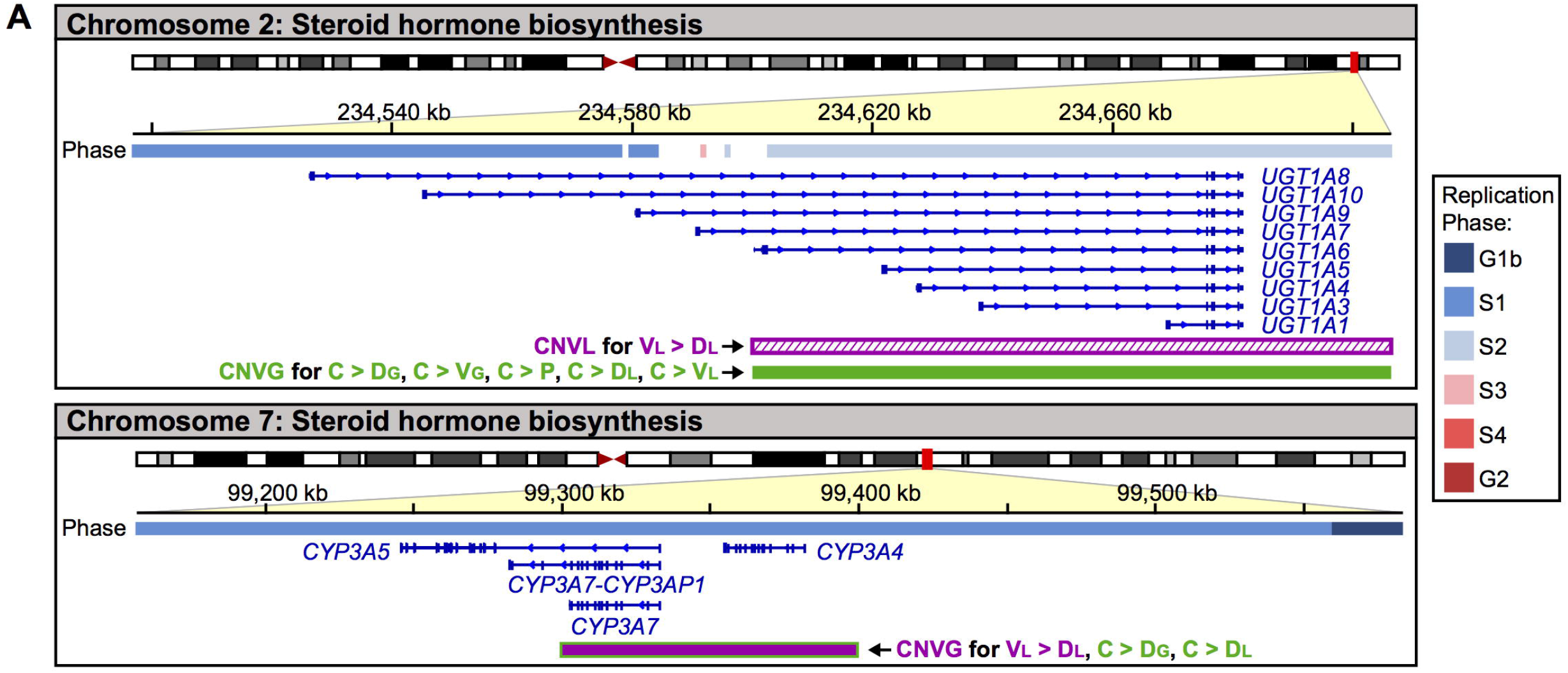
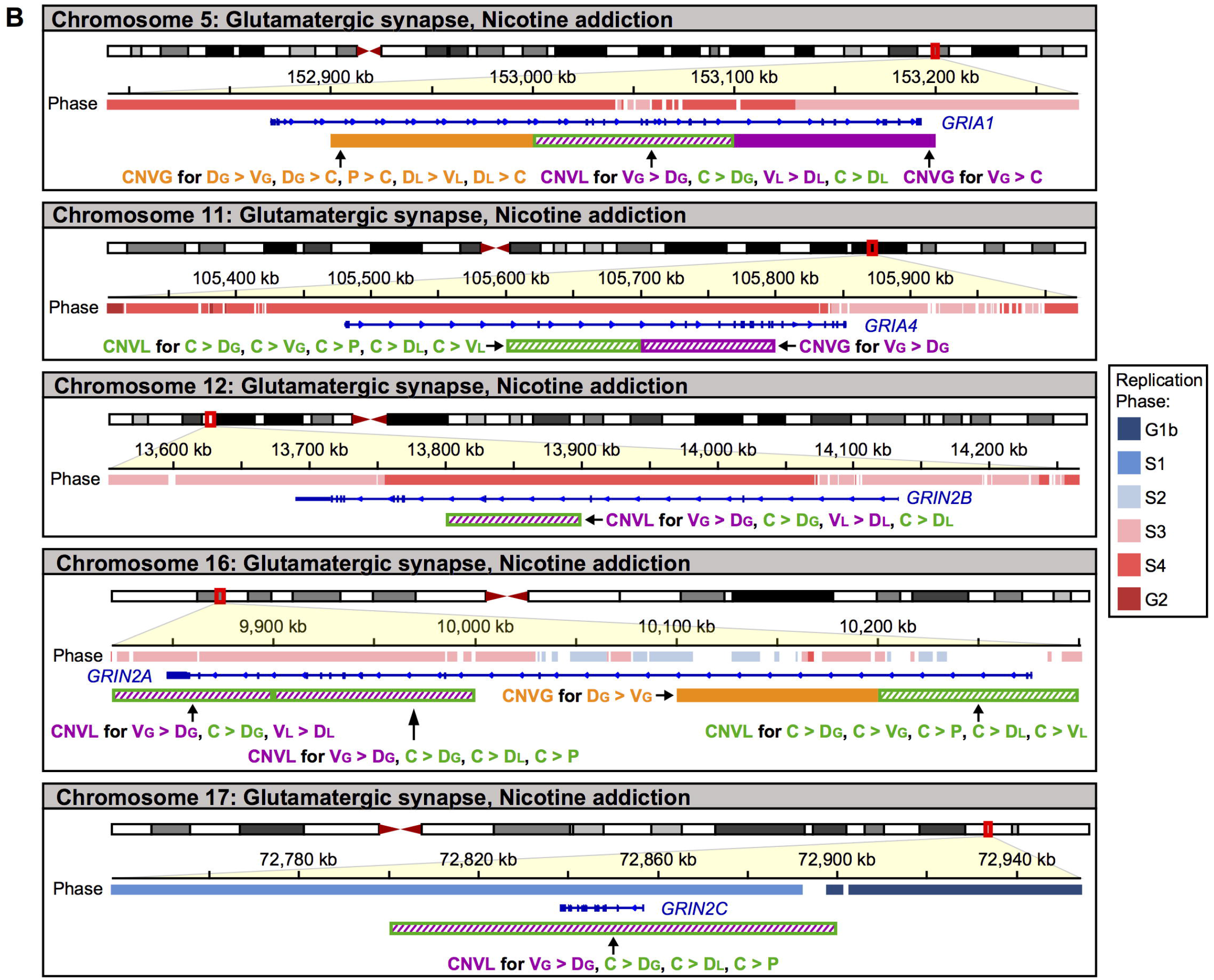
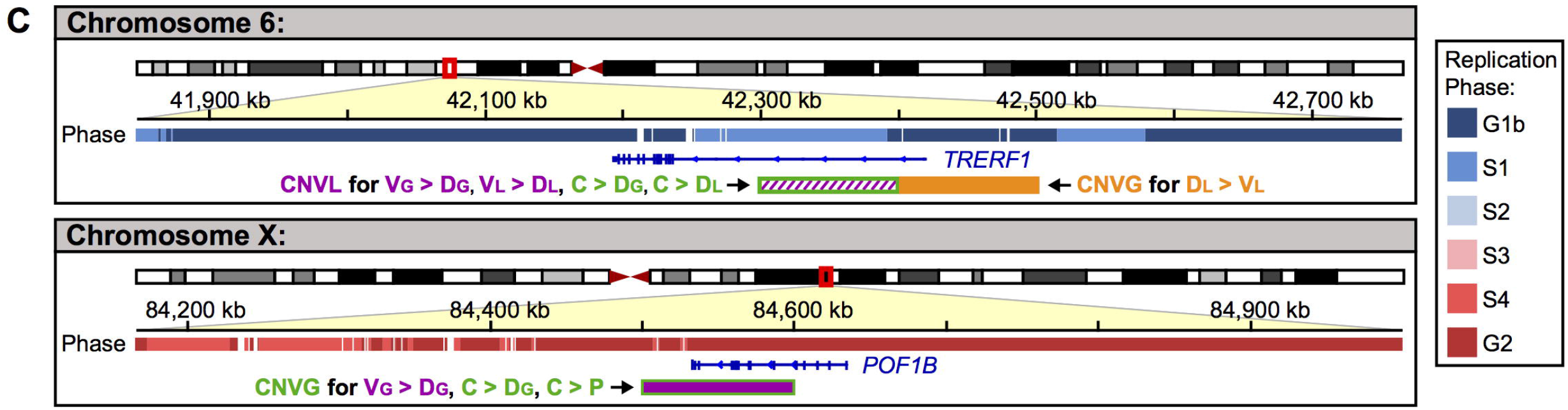
Selected genes overlapping with frequency-based diagnostic CNV features. Expanded views of chromosomal segments on (A) chromosomes 2 and 7 for steroid biosynthesis pathway genes, (B) chromosomes 5, 11, 12, 16 and 17 for *GRI*-genes of the glutamatergic synapse and nicotine addiction pathways, and (C) chromosomes 6 and X for the non-pathway *TRERF1* and *POF1B* genes with color-coded representation of the DNA replication phase in the ‘Phase’ track, and aligned gene sequence(s) in blue (e.g. *UGT1A8* or *TRERF1*) as described in RefSeq Genes in UCSC Genome Browser. Green rectangular boxes either below the genes indicate the presence of diagnostic CNVG or CNVL feature(s). Inside each box, colored stripes are indicative of CNVL features(s), and solid coloring is indicative of CNVG features(s): purple for predominantly V-favoring features, orange for D-favoring features, and green for C-favoring features.

### Pathways and genes enriched in diagnostic CNV features

A wide range of genes showed sequence overlaps with the frequency-based diagnostic CNVG and CNVL features of a range of KEGG pathways in PMDD and its subtypes (Table 1) which pointed to their possible contributions to the PMDD disorder, and some major genes were contained in more than one pathway (Table 2). It was striking that, as indicated in lines 1-5 of Table 2, the control C-type was favored by high frequencies of CNVG features relative to the diseased P-, D- or V-type, suggesting that a major causal factor of the PMDD disorder could be decreased levels of the CNVG features overlapping with the steroid hormone biosynthesis pathway, with the involvement of *CYP-* and *UGT-genes* replicating in phases S2 and S1. As shown in lines 9-17 of Table 2, the C-type and V-type profiles were favored over the D-type by high frequencies of CNVL features in the *GRI-genes*, which were involved in pathways of nicotine addiction, circadian entrainment, serotonergic synapse, dopaminergic synapse and cAMP signaling. The chromosomal sites of these genes and their overlaps with the 100-kb CNV features are shown in Figure 5.

**Table 1.**
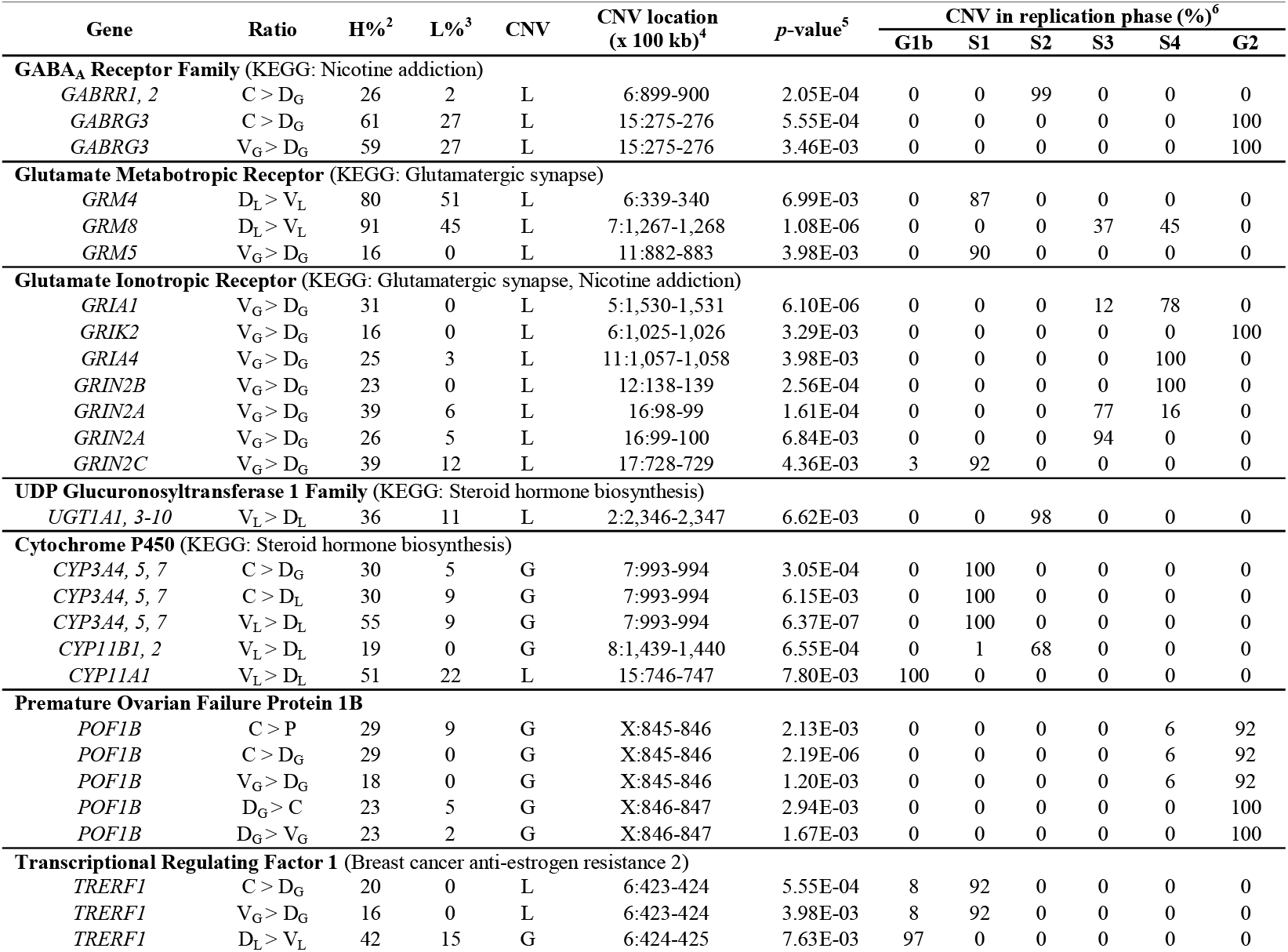
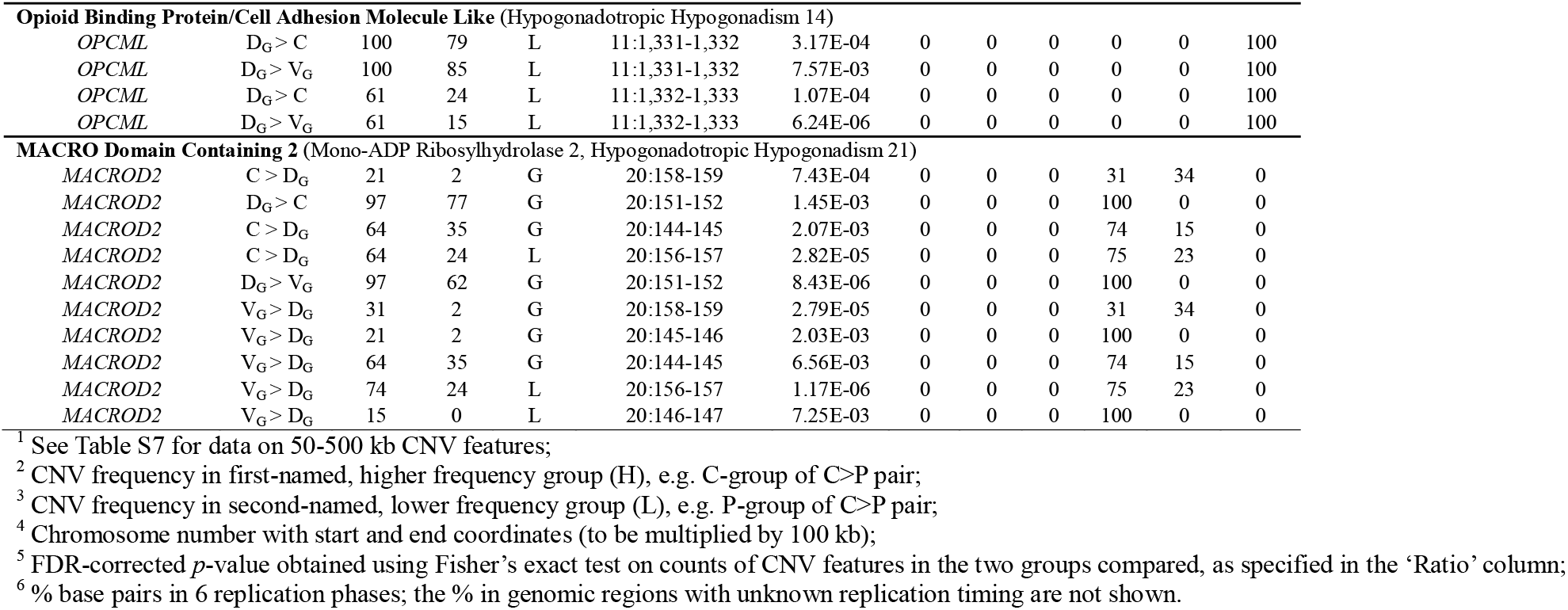
Selected genes overlapping with 100-kb frequency-based CNVG and CNVL features^1^ with adjusted *p*-values less than 0.01.

**Table 2.**
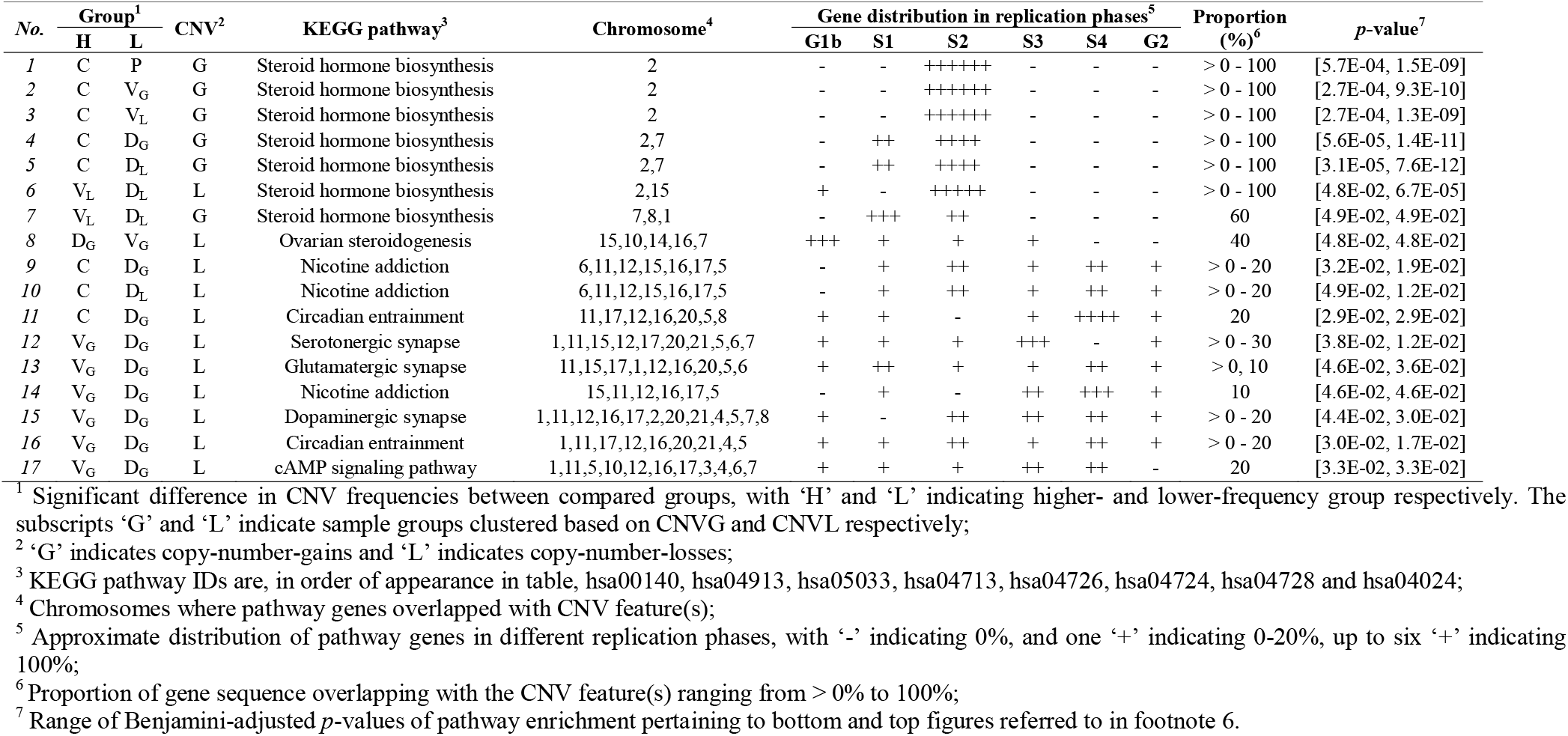
Representative pathways enriched in 100-kb frequency-based CNV features.

The 50-kb CNV features overlapped with the genes in the glutamatergic-synapse, alcoholism, and systemic lupus erythematosus pathway genes, as well as steroid hormone biosynthesis pathway genes replicating in S2 and S3 (Table S5). On the other hand, the 450-kb CNV features overlapped with chemokine signaling pathway genes (Table S5). Because high regional density of genes could impact on gene annotations by yielding false-positive co-localizations when a CNV feature incidentally captured a gene cluster belonging to a pathway, empirical *p*-values based on Monte Carlo simulations were also estimated for the 100-kb CNV features (Table S10), which provided additional support for some of the pathway in Table 2 through the elimination of such false positives (see ‘Statistical analysis’ in Methods).

### Genomic features enriched in diagnostic CNV features

Co-localization analysis revealed various associations between 100-kb frequency-based CNV features and a wide spectrum of genomic features in different replication phases (Table 3 and Table S11). D versus V differences in genomic feature contents can be identified from the thermal scale plots of co-localization scores illustrated in Figure 6 and Figure S8. The genomics features apparently differed between D and V types included: (1) In terms of retrotransposons, D-favoring CNVG features enriched with more of the subfamily of evolutionarily very young short transposons SVAef, while V-favoring CNVG and D-favoring CNVL features enriched with the very young long transposon subfamily, L1vy. (2) With respect to genetic markers, P-favoring, especially D-favoring CNVL features were enriched with recombination events as well as genetic variation hotspots and clusters (27). GWAS reported markers were co-localized with D-favoring CNVGs in S1, V-favoring CNVLs in S4 and C-favoring CNVLs G1b. As well, ClinVar markers were enriched in V-favoring CNVG of S2 phase and D-favoring CNVL of S1. (3) In respect to the group of CpG-related genomic features, the main difference between the two types was that D-favoring CNVL features were more enriched with CpG features such as MeBS in S4 replicating sequences. Compared with D- and V-favoring, the C-favoring CNV features were more prominently enriched with CpG features, especially for C-favoring CNVG in S3 and CNVL in G2 and S4 phases. (4) In regard to non-coding RNA, LINC was enriched in V-favoring CNVL as well as C-favoring CNV features, but not in D-favoring features. (5) To a lesser extent, the enrichment of histone binding sites in D-favoring CNVG features of G2 and S2 phases. In contrast, histone sites were enriched in V-favoring CNVL features of S4 phase. This trend was clearly visible from Figure 6, where twelve kind histone binding sites were analyzed separately and displayed side-by-side.

**Table 3.** Selected genomic features in 100-kb frequency-based diagnostic CNV features with fold-change greater than 1 in at least one replication phase(s).

| Group<sup>1</sup> |  |  | Fold-change in different replication phases<sup>3</sup> |  |  |  |  |  |
| --- | --- | --- | --- | --- | --- | --- | --- | --- |
| H | L | CNV<sup>2</sup> | G1b | S1 | S2 | S3 | S4 | G2 |
| <u>**CpGi (CpG island)**</u> |  |  |  |  |  |  |  |  |
| C | P | G | -0.32 | 0.06 | -0.06 | **1.62** | -0.18 | **1.15** |
| C | P | L | -0.18 | 0.05 | 0.39 | 0.46 | **1.10** | **2.20** |
| C | D<sub>L</sub> | G | -0.45 | 0.02 | -0.08 | 0.59 | **1.18** | 0.57 |
| C | V<sub>G</sub> | L | 0.14 | 0.06 | 0.69 | 0.71 | 0.47 | **1.42** |
| P | C | L | -0.34 | 0.12 | -0.02 | 0.38 | -0.23 | **1.08** |
| <u>**Me450 (Methylation status using HumanMethylation450)**</u> |  |  |  |  |  |  |  |  |
| C | P | G | -0.51 | 0.54 | **1.04** | 0.88 | -0.01 | 0.43 |
| C | P | L | -0.29 | 0.23 | -0.02 | 0.33 | **1.16** | **1.05** |
| C | V<sub>L</sub> | G | **1.00** | 0.04 | -0.02 | -0.35 | -0.48 | -0.25 |
| <u>**MeBS (cytosine methylation using bisulfite sequencing)**</u> |  |  |  |  |  |  |  |  |
| C | P | L | -0.12 | 0.45 | 0.12 | 0.49 | **1.53** | **1.55** |
| C | D<sub>L</sub> | G | -0.53 | 0.10 | 0.06 | 0.64 | **1.65** | 0.59 |
| C | V<sub>G</sub> | L | 0.27 | 0.26 | 0.69 | 0.94 | **1.51** | 0.29 |
| D<sub>G</sub> | C | L | **1.03** | 0.54 | 0.35 | 0.24 | **1.20** | -0.39 |
| D<sub>L</sub> | C | L | **1.38** | 0.20 | 0.25 | 0.37 | 0.13 | -0.34 |
| D<sub>G</sub> | V<sub>G</sub> | L | **1.15** | 0.41 | 0.37 | 0.46 | **1.69** | -0.24 |
| D<sub>L</sub> | V<sub>L</sub> | L | 0.82 | 0.45 | 0.53 | 0.59 | **1.47** | -0.15 |
| V<sub>L</sub> | D<sub>L</sub> | G | 0.15 | -0.17 | -0.14 | 0.45 | **1.35** | **1.21** |
| <u>**MeMRE (Methylation using MRE-**</u> |  |  |  |  |  |  |  |  |
| C | P | G | -0.26 | 0.23 | -0.30 | **2.00** | -0.12 | -0.19 |
| C | P | L | -0.23 | -0.06 | 0.31 | 0.47 | **1.44** | 0.90 |
| P | C | L | -0.13 | 0.10 | 0.10 | 0.33 | -0.22 | **1.70** |
| <u>**DNase (DNase I hypersensitive sites)**</u> |  |  |  |  |  |  |  |  |
| C | P | L | -0.12 | 0.02 | 0.11 | 0.55 | 0.46 | **1.02** |
| C | V<sub>L</sub> | G | 0.22 | -0.30 | 0.25 | -0.55 | 0.04 | **1.14** |
| P | C | L | -0.10 | 0.08 | 0.51 | 0.38 | **1.80** | -0.13 |
| D<sub>G</sub> | C | L | 0.28 | 0.27 | 0.24 | -0.05 | **1.37** | -0.52 |
| D<sub>L</sub> | C | L | 0.47 | 0.22 | 0.39 | -0.12 | **1.77** | -0.14 |
| V<sub>G</sub> | C | L | -0.16 | 0.22 | 0.05 | 0.00 | **1.38** | -0.57 |
| V<sub>L</sub> | C | L | -0.15 | 0.14 | 0.25 | 0.09 | **1.08** | -0.66 |
| <u>**FAIRE (Formaldehyde-assisted isolation of regulatory elements)**</u> |  |  |  |  |  |  |  |  |
| C | P | L | -0.18 | 0.17 | -0.10 | 0.15 | 0.71 | **2.69** |
| C | V<sub>G</sub> | L | -0.23 | 0.12 | 0.03 | 0.14 | 0.46 | **2.97** |
| C | V<sub>L</sub> | L | 0.02 | 0.08 | -0.04 | 0.04 | 0.33 | **2.54** |
| P | C | L | -0.09 | 0.16 | 0.53 | -0.03 | 0.86 | **1.56** |
| D<sub>G</sub> | C | L | 0.32 | 0.28 | 0.16 | -0.17 | **1.55** | 0.14 |
| D<sub>L</sub> | C | L | 0.34 | 0.29 | 0.33 | -0.24 | **2.11** | **1.25** |
| D<sub>G</sub> | V<sub>G</sub> | L | 0.22 | 0.17 | 0.11 | 0.11 | **1.15** | 0.92 |
| D<sub>L</sub> | V<sub>L</sub> | L | 0.20 | 0.21 | 0.02 | 0.03 | 0.77 | **2.08** |
| H | L | CNV<sup>2</sup> | G1b | S1 | S2 | S3 | S4 | G2 |
| <u>**LINC (Large intergenic non-coding RNA)**</u> |  |  |  |  |  |  |  |  |
| <u>**Continued**</u> |  |  |  |  |  |  |  |  |
| C | P | G | -0.97 | -0.52 | **18.30** | **4.23** | 0.32 | -0.11 |
| C | P | L | -0.99 | **1.05** | 0.17 | **1.02** | -0.73 | 0.92 |
| C | V<sub>L</sub> | G | 0.11 | -0.96 | 0.03 | **1.75** | -0.55 | **2.42** |
| C | V<sub>G</sub> | L | -1.00 | -0.77 | -0.79 | -0.93 | -0.47 | **1.59** |
| C | V<sub>L</sub> | L | -0.98 | -0.74 | -0.72 | -0.87 | -0.50 | **1.06** |
| V<sub>G</sub> | C | L | 0.48 | -0.70 | 0.32 | **2.14** | **1.61** | **1.36** |
| V<sub>L</sub> | C | L | 0.31 | -0.75 | -0.02 | **1.80** | **1.01** | 0.74 |
| V<sub>G</sub> | D<sub>G</sub> | L | 0.37 | -0.67 | **1.19** | 0.52 | 0.33 | **1.34** |
| <u>**RecD (Sex-averaged rates of recombination)**</u> |  |  |  |  |  |  |  |  |
| P | C | L | **1.31** | 0.39 | 0.26 | 0.44 | 0.36 | 0.97 |
| D<sub>G</sub> | C | L | 0.23 | 0.16 | 0.43 | 0.42 | 0.35 | **1.24** |
| D<sub>L</sub> | C | L | 0.40 | 0.30 | 0.52 | 0.50 | 0.51 | **1.56** |
| D<sub>L</sub> | V<sub>L</sub> | L | 0.25 | 0.13 | 0.24 | 0.22 | 0.45 | **1.06** |
| <u>**GWAS (GWAS-identified SNPs)**</u> |  |  |  |  |  |  |  |  |
| C | V<sub>G</sub> | L | **2.08** | -0.01 | -0.02 | -0.35 | 0.18 | 0.60 |
| C | V<sub>L</sub> | L | **1.63** | -0.13 | -0.01 | -0.23 | 0.10 | 0.38 |
| P | C | G | -0.26 | **5.48** | -0.38 | 0.27 | -0.26 | -0.34 |
| D<sub>G</sub> | C | G | -0.16 | **1.61** | -0.13 | -0.28 | -0.05 | -0.01 |
| D<sub>L</sub> | C | G | -0.08 | **1.78** | -0.17 | -0.22 | -0.03 | -0.16 |
| V<sub>G</sub> | C | L | -0.34 | -0.20 | -0.22 | -0.23 | **2.78** | -0.14 |
| V<sub>L</sub> | C | L | -0.28 | -0.37 | -0.29 | -0.17 | **2.00** | 0.01 |
| D<sub>G</sub> | V<sub>G</sub> | G | -0.16 | **1.25** | -0.02 | 0.33 | -0.28 | -0.25 |
| D<sub>L</sub> | V<sub>L</sub> | G | -0.08 | **1.27** | -0.03 | 0.24 | -0.24 | -0.11 |
| <u>**GV hotspot (Density-based genetic-variant hotspot)**</u> |  |  |  |  |  |  |  |  |
| C | V<sub>G</sub> | G | **1.97** | -0.34 | -0.21 | -0.27 | -0.15 | **1.22** |
| C | V<sub>L</sub> | G | **1.63** | 0.01 | -0.27 | -0.09 | -0.28 | 0.69 |
| P | C | L | 0.12 | -0.01 | -0.11 | 0.70 | **1.58** | 0.29 |
| D<sub>G</sub> | V<sub>G</sub> | G | -0.03 | -0.23 | -0.44 | 0.14 | 0.20 | **1.03** |
| D<sub>L</sub> | V<sub>L</sub> | G | 0.01 | -0.31 | -0.43 | 0.28 | 0.00 | **1.06** |
| D<sub>G</sub> | V<sub>G</sub> | L | 0.30 | 0.27 | 0.09 | 0.42 | 0.61 | **1.22** |
| <u>**Cluster (Cluster of genetic-variant hotspots)**</u> |  |  |  |  |  |  |  |  |
| C | D<sub>L</sub> | L | 0.03 | -0.84 | 0.38 | -0.87 | -0.30 | **2.42** |
| C | V<sub>G</sub> | G | **3.68** | -1.00 | -1.00 | 0.42 | -0.15 | -0.29 |
| C | V<sub>L</sub> | G | **2.90** | -1.00 | -1.00 | 0.37 | -0.21 | -0.52 |
| C | V<sub>G</sub> | L | -0.23 | **1.42** | 0.03 | 0.24 | 0.26 | 0.83 |
| P | C | L | -1.00 | -1.00 | -0.27 | **4.33** | **4.26** | -1.00 |
| D<sub>G</sub> | C | L | **2.15** | 0.73 | -0.05 | 0.57 | **1.34** | -1.00 |
| D<sub>L</sub> | C | L | **2.46** | 0.90 | 0.07 | 0.73 | **1.11** | -1.00 |

| <u>*LINC (Large intergenic non-coding RNA)*</u> |  |  |  |  |  |  |  |  |  |  |  |  |  |  |  |  |  |
| --- | --- | --- | --- | --- | --- | --- | --- | --- | --- | --- | --- | --- | --- | --- | --- | --- | --- |
| C | D<sub>G</sub> | G | 0.50 | -0.64 | **7.56** | **1.38** | **1.46** | 0.11 | V<sub>G</sub> | C | G | -1.00 | -0.26 | **1.48** | 0.46 | 0.12 | -0.83 |
| C | D<sub>L</sub> | G | -0.98 | -0.76 | **8.90** | **1.53** | **1.51** | 0.16 | V<sub>L</sub> | C | G | -1.00 | **1.45** | 0.72 | 0.19 | -0.08 | -0.38 |
| C | D<sub>G</sub> | L | -0.86 | 0.44 | 0.46 | -0.79 | -0.88 | **3.10** | V<sub>G</sub> | C | L | -0.41 | -0.72 | -0.70 | **1.68** | 0.46 | -0.97 |
| D<sub>G</sub> | C | G | **1.84** | -0.92 | -0.68 | 0.21 | -0.99 | -0.80 | V<sub>L</sub> | C | L | -0.21 | -0.78 | -0.41 | **1.20** | 0.27 | -0.89 |
| D<sub>G</sub> | C | L | -0.85 | -0.84 | -0.95 | **1.89** | -1.00 | -1.00 | D<sub>G</sub> | V<sub>G</sub> | G | 0.02 | -0.65 | -0.55 | -0.50 | **1.52** | 0.89 |
| D<sub>L</sub> | C | L | -0.81 | -0.82 | -0.95 | **1.85** | -1.00 | -1.00 | D<sub>L</sub> | V<sub>L</sub> | G | -0.05 | -0.64 | -0.55 | -0.53 | **1.41** | 0.95 |
| V<sub>G</sub> | D<sub>G</sub> | G | 0.87 | -0.74 | **4.34** | -0.66 | 0.81 | 0.16 | D<sub>G</sub> | V<sub>G</sub> | L | 0.91 | 0.42 | 0.43 | 0.64 | **1.71** | **1.25** |
| V<sub>L</sub> | D<sub>L</sub> | G | 0.76 | -0.75 | **3.45** | -0.69 | 0.65 | -0.08 | V<sub>G</sub> | D<sub>G</sub> | L | -0.64 | -0.36 | **3.00** | 0.50 | 0.12 | 0.20 |
|  |  |  |  |  |  |  |  |  | V<sub>L</sub> | D<sub>L</sub> | L | -0.44 | -0.52 | -0.15 | 0.41 | -0.38 | **1.07** |

<sup>1</sup> Significant difference in CNV frequencies between compared groups, with 'H' and 'L' indicating higher- and lower-frequency group respectively. The subscripts 'G' and 'L' indicate sample groups clustered based on CNVG and CNVL respectively;
<sup>2</sup> 'G' indicates copy-number-gains and 'L' indicates copy-number-losses;
<sup>3</sup> Fold-change (> 1-fold in bold) of genomic feature density or intensity in diagnostic CNV features relative to non-diagnostic-CNV regions in replication phase.

**Figure 6.**
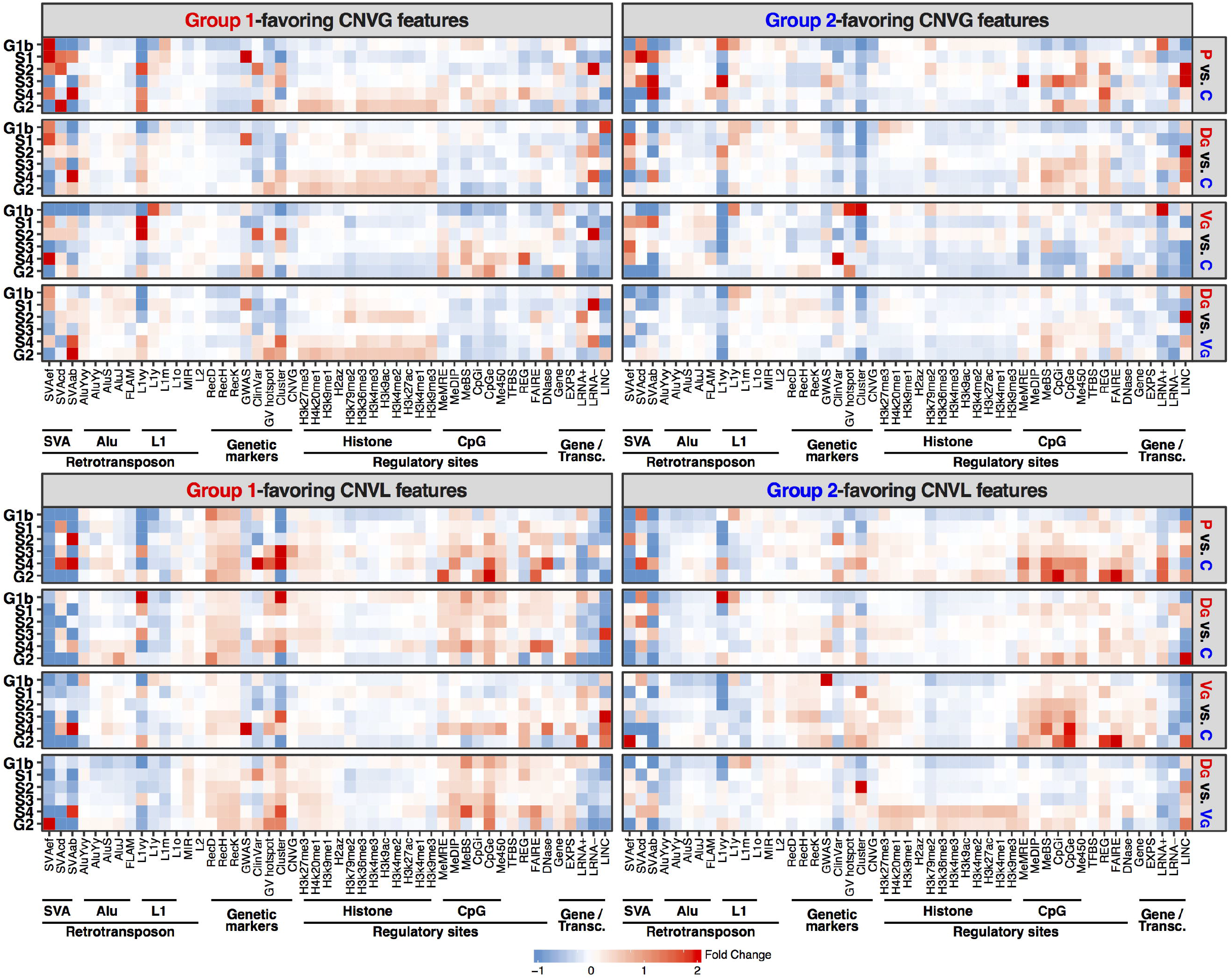
Enrichment analysis of genomic-feature contents in different replication phases for control and PMDD subtypes. Frequency-based CNV features diagnostic for C group, i.e., control, as well as that for D and C groups of PMDD samples clustered by CNVG dendrogram, identified with 100-kb scanning windows, were used in the analysis. Enrichment analysis results were plotted for CNVG features in the upper two panels and that for CNVL features in the bottom two panels. A similar analysis performed in parallel for clustered by CNVL dendrogram can be found in Figure S8. Fold-change of each genomic feature in the diagnostic CNV features relative to the non-diagnostic-CNV regions was estimated according to ‘Genomic-feature content of diagnostic CNV features in different replication phases’ in Methods, and was color-coded based on the thermal scale. Fold-change greater than 2-fold was capped at 2 in the heat map. ‘Group 1’ indicated the first-named group and ‘Group 2’ the second-named group in a given pair of samples. Genomic features were grouped into Retrotransposon (SVAef, SVAcd, SVAab, AluYvy, AluYy,s AluS, AluJ, FLAM, L1vy, L1y, L1m, L1o, MIR, L2), Genetic markers (RecD, RecH, RecK, GWAS, ClinVar, GV hotspot, Cluster, CNVG), Regulatory sites (H3k27me3, H4k20me1, H3k9me1, H2az, H3k79me2, H3k36me3, H3k4me3, H3k9ac, H3k4me2, H3k27ac, H3k4me1, H3k9me3, MeMRE, MeDIP, MeBS, CpGi, CpGe, Me450, TFBS, REG, FAIRE, DNase) and Gene/Transcription (Gene, EXPS, LRNA+, LRNA-, LINC) groups on the x-axis based on their sequence and functional properties. The descriptions of genomic features and numeric data were available in Table S11.

Some of the strongly enriched genomic features with great than one-fold enrichment was listed in Table 3. For example, enrichment of DNase I hypersensitive sites (DNase) was found in C>P CNVL and C>V_L_ CNVG features in G2-phase replicating sequences and P>C (as well as D>C and V>C) CNVL features in S4-phase replicating sequences. Regulatory elements isolated by formaldehyde (FAIRE) were found to enrich in C>V, D>C and D>V CNVL features that located in the late-replicating S4 and G2 phases. Disease- or trait-associated SNPs identified by genome-wide association studies (GWAS) were enriched in C>V CNVLs in G1b phase, and P>C CNVGs in S1 phase reaching a fold-change of 5.5 relative to non-diagnostic-CNV regions in S1 phase. The C-favoring (C>P and C>D_L_) CNVG features and C-favoring CNVL (C>P and C>V_G_) features tend to co-localize with CpG islands (CpGi) in median to late-replicating S3-G2 phases. A range of methylation-related features (Me450, MeBS, and MeMRE) were found to enrich in C-favoring CNVG features mainly in early to median G1b-S3 phases, and C-favoring CNVL features in late-replicating S4-G2 phases. Long intergenic non-coding RNAs (LINC) were found to be enriched in C-favoring CNVGs mainly in S2 phase or CNVLs mainly in G2 phase, D-favoring CNVGs in G1b phase, and V-favoring CNVLs in S3-G2 phases.

## Discussion

Application of either the cutree method or the semi-supervised method to the hierarchically clustered CNVGs or CNVLs in the germline genomes of PMDD subjects enabled the distinction between the D-type and V-type CNV profiles. The high degree of consistency between the clinical depression-subtype and D-type CNV profiles, and between the clinical invasion-subtype and V-type CNV profiles, indicated that the two clinical PMDD subtypes were intrinsically correlated with the two dissimilar types of CNV profiles. This was further conformed when diagnostic CNVG and CNVL features were selected by means of machine learning using the correlation method, and employed as abundance markers to predict whether a given germline genomic sample belonged to the control group, the PMDD group, the V-type CNV group or the D-type CNV group, yielding average accuracies of prediction of 81.0-88.4% (Figure 3), which in turn validated the use of diagnostic CNVG and CNVL features to identify the genes and pathways that overlapped with such diagnostic features as potential contributors to the PMDD disorder.

In this regard, there exists overall accord between the cutree and the semi-supervised methods in terms of diagnostic CNV features identified, replication-phase distribution and pathway enrichments (Table S5, S12 and S13). As indicated in DSM-V, PMDD is defined by a complex system of symptoms. In the present study, limited data allowed the analysis of only the depression-type and invasion-type symptoms. Nevertheless, the Venn diagrams in Figure 3 clearly showed that the CNVs underlying the D-type and V-type CNV profiles were strikingly more divergent from one another than their separate divergences from the CNVs underlying the C-type. This finding was also consistent with the results in Figure 2A, which showed that there were more correlation-based or frequency-based CNV features that could be employed to distinguish between D_G_-vs-V_G_ or D_L_-vs-V_L_ compared to CNV-features that could distinguish between D_G_-vs-C, V_G_-vs-C, D_L_-vs-C or V_L_-vs-C. As well, the mixed distribution of control CNV profiles among depression- or invasion-type CNV profiles (Figure S3) indicated that the difference between the CNVs in the two subtypes of PMDD was larger than their individual differences from the control. This surprising genome condition, as illustrated in Figure 7, raises the question of whether the depression-type and invasion-type conditions of PMDD might represent two distinct clinical disorders.

**Figure 7.**
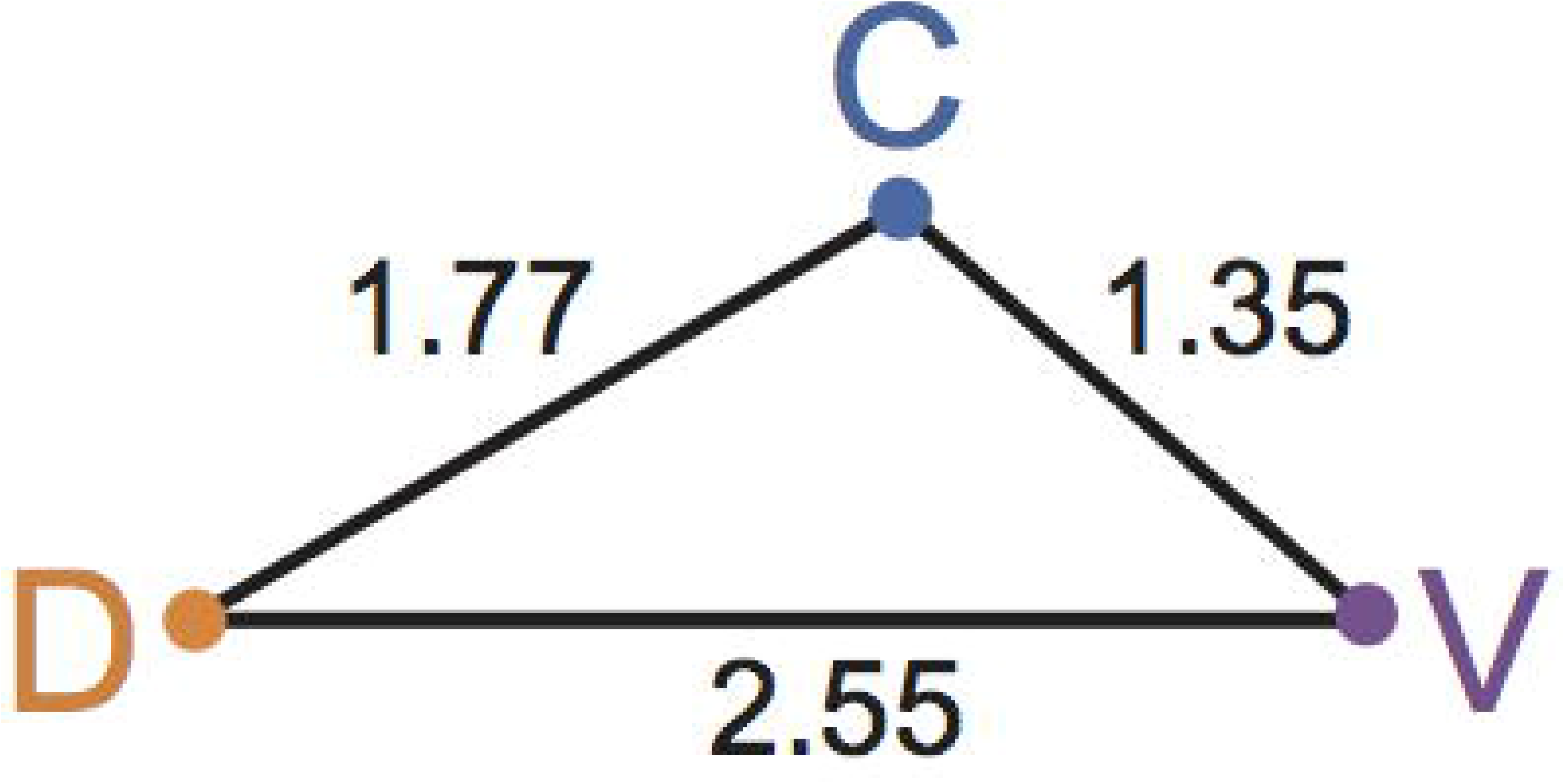
Genetic distances between the two subtypes of PMDD and the control. Pairwise distances were estimated based on the abundance of diagnostic CNV features between C-, D- and V-type genomes. The numbers of frequency-based CNV features were employed as an approximate index of the genetic distance between the D-vs-C, V-vs-C or D-vs-V sample pairs in Table S15, which comprised the 50-500 kb frequency-based CNV features. Notably, the D-vs-V distance was larger than the D-vs-C distance or the V-vs-C distance.

A faithful temporal order of DNA replication is fundamental to normal cellular function, and aberrant replication timings were observed in complex diseases including cancers (34, 35). Accordingly, the relative abundances of diagnostic CNVG and CNVL features among genomic DNA sequence regions preferentially replicating in each one of the six phases of cell cycle, namely G1b, S1, S2, S3, S4 and G2 were examined in Figure 4. The peaks of C-favoring CNVL features in P-vs-C comparisons (downward hollow green bars in Figure 4A) shifted clearly from the early S1 phase among the 50-kb features to the late G2 phase among the 450-kb features, pointing to the enrichment of some small CNVL features in the early-replicating regions and larger CNVL features in the late-replicating regions among the determinants of the C-type, *viz*. in the prevention of PMDD occurrence. As well, more D-favoring CNVG features were located in G2-replicating sequences compared to genomic DNA sequences replicating in other cell-cycle phases within the D>C and D>V groups, which was particularly notable in view of the enrichment of G2 phase-replicating sequences in non-coding sequences (27, 29). In addition, the abundance of V-favoring 50-kb CNVL features in the V>C and V>D comparisons (Figure 4B and D) suggests that small-size CNVLs also played important roles in the development of V-type PMDD.

When diagnostic CNVs were analyzed for their genomic feature enrichment with reference to replication phases, interesting observations were obtained (Figure 6; Table 3). It has been revealed that the late-replicating S4-G2 phases in the gene-distal zones are found to be depleted of functional genomic features (27). However, the present study observed associations of open chromatin signals, regulatory elements and epigenetic regulation sites with the diagnostic CNV features in these late-replicating sequences (Table 3), indicating that the diagnostic CNV features might represent pivotal genomic sites in the late-replicating sequences that sequence alterations may give raise to functional perturbations underlying PMDD and its two subtypes. As illustrated herein, genomic feature content analysis, implemented with replication phase information, has pointed to the likelihood of genomic events underlying the subtyping of PMDD and hence a genomic nature of the disorder and its clinical diversity. Since genomic features included in the analysis were broad in spectrum and well beyond the boundary of known genes, the feature enrichment analysis performed herein may complement with and surpass genetic pathway analysis as a powerful tool for genomic studies on complex traits and disorders.

Previously, a number of genes was proposed to be PMDD suspectable genes, including those of steroid hormone biosynthesis (2, 3), and estrogen signaling (36, 37), and these proposals were supported by the presence of these genes in Table 1. The overlaps of genes of nicotine addiction, glutamatergic synapses, olfactory transduction, alcoholism, systemic lupus erythematosus, hypogonadism, premature ovarian failure, and breast cancer with PMDD might be suggestive of hitherto hidden aspects of central nervous system or endocrine system involvements with PMDD. The *GRIA4* gene, overlapping with the 100-kb CNV features for the C>D and V>D comparisons, groups, has also been found to be associated with schizophrenia (38), in accordance with the shared CNVs between schizophrenia and PMDD (14).

In conclusion, through CNV profiling, the present study provided evidence for strong correlation of the clinical depression-subtype or invasion-subtype with the D-type and V-type germline genomes, marked by the overlaps between their CNVs and the machine-selected diagnostic CNV features that favored one or another type of genomes. On account of this correlation, the diagnostic CNV features could be employed as frequency markers to predict the propensity to PMDD and one of its clinical subtypes, as well as position markers to identify candidate PMDD genes and pathways. Moreover, the genetic difference between the depression-favoring and invasion-favoring CNV profiles was found to exceed their individual divergences from the normal controls (Figure 7), raising the question of how this outcome might have been evolved. Future studies will be required to determine how many of the array of PMDD symptoms besides the depression-subtype and invasion-subtype ones could be significantly correlated with CNVs, and what complex diseases other than PMDD would embody CNV-symptom correlations as strong as those encountered with PMDD.

## Acknowledgements

We thank Ms. Peggy Lee for the technical support. The study was supported by grants to HX from University Grants Council (SRF116SC01; UROP18SC06; UROP20SC07) and Innovation Technology Council (ITS/085/10; ITS113/15FP; ITCPD/17-9; ITT/023/17GP; ITT/026/18GP) of Hong Kong SAR; Shenzhen Municipal Council of Science and Technology, Guangdong (JCYJ20170818113656988); Shandong Province First Class Disciple Development Grant and Tai-Shan Scholar Program, Shandong; and Ministry of Science and Technology (National Science and Technology Major Project, No. 2017ZX09301064), People’s Republic of China, as well as grants from National Natural Science Foundation of China to MQ (No. 8157151623) and JW (No. 81603510), respectively.

## Author Contributions

HX and MQ conceived and designed the experiments, ZW, XLo, AU, SC and WM performed the AluScan sequencing related experiments and analysis of the sequencing data. PS, MG, JW, HW, XLi, WS and MQ coordinated the collection of PMDD and control cohorts, and HX, ZW, XLo, SC and MQ wrote the paper.

## Supplementary Information

Supplementary materials are available online, including Figures S1-S8 and Table S1-S15.

