## Supplementary Figures S1-S8 for "Copy number variation profile-based genomic subtyping of premenstrual dysphoric disorder in Chinese"

#### Contents:

- Supplementary Figures S1 to S8
- Caption of Supplementary Tables S1 to S15

### Supplementary Figures

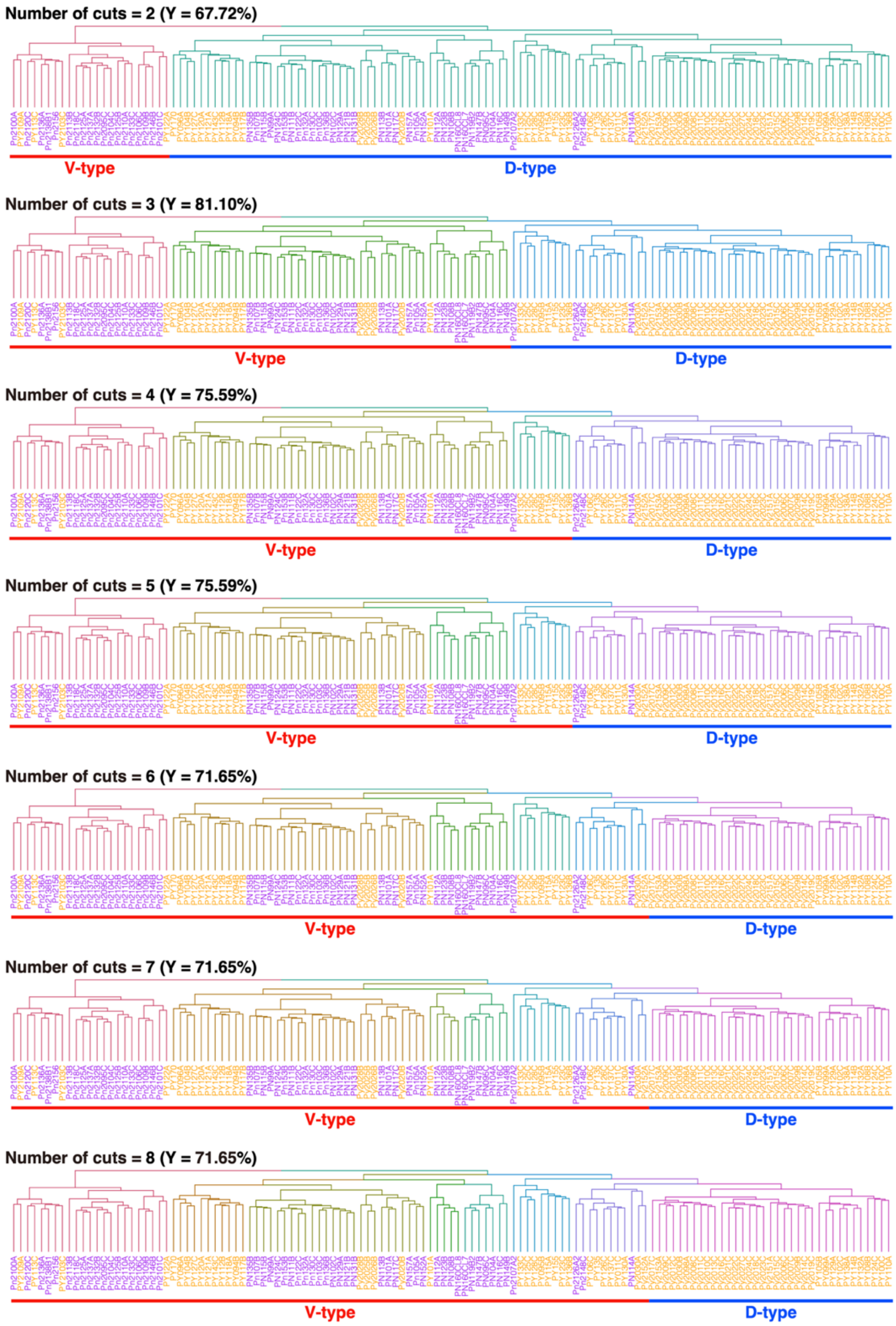


**Figure S1. Division of 100-kb CNVG-profile dendrogram into 2 to 8 sub-clusters.** Branches of distinct cuts on the dendrogram obtained using the cutree method are shown in different colors. The samples with IDs in purple represent the clinical depression-subtype, and the samples with IDs in orange represent the clinical invasion-subtype. On the other hand, all the samples underlined by blue line with 'D-type' label are designated as genomes with D-type CNV profiles, and all the remaining samples underlined by red line with 'V-type' label are designated as genomes with V-type CNV profiles. Upon comparison of the different number of cuts, the results showed that CNV-based subtyping of D- and V-types with cut number = 3 yielded the highest level of consistency (Y) with clinical subtyping in accordance with Eqn 2. Hence the D- and V-types delineated using cut number = 3 were employed as the optimal cutree CNVG profiles for the analysis of 100 kb-CNVGs.

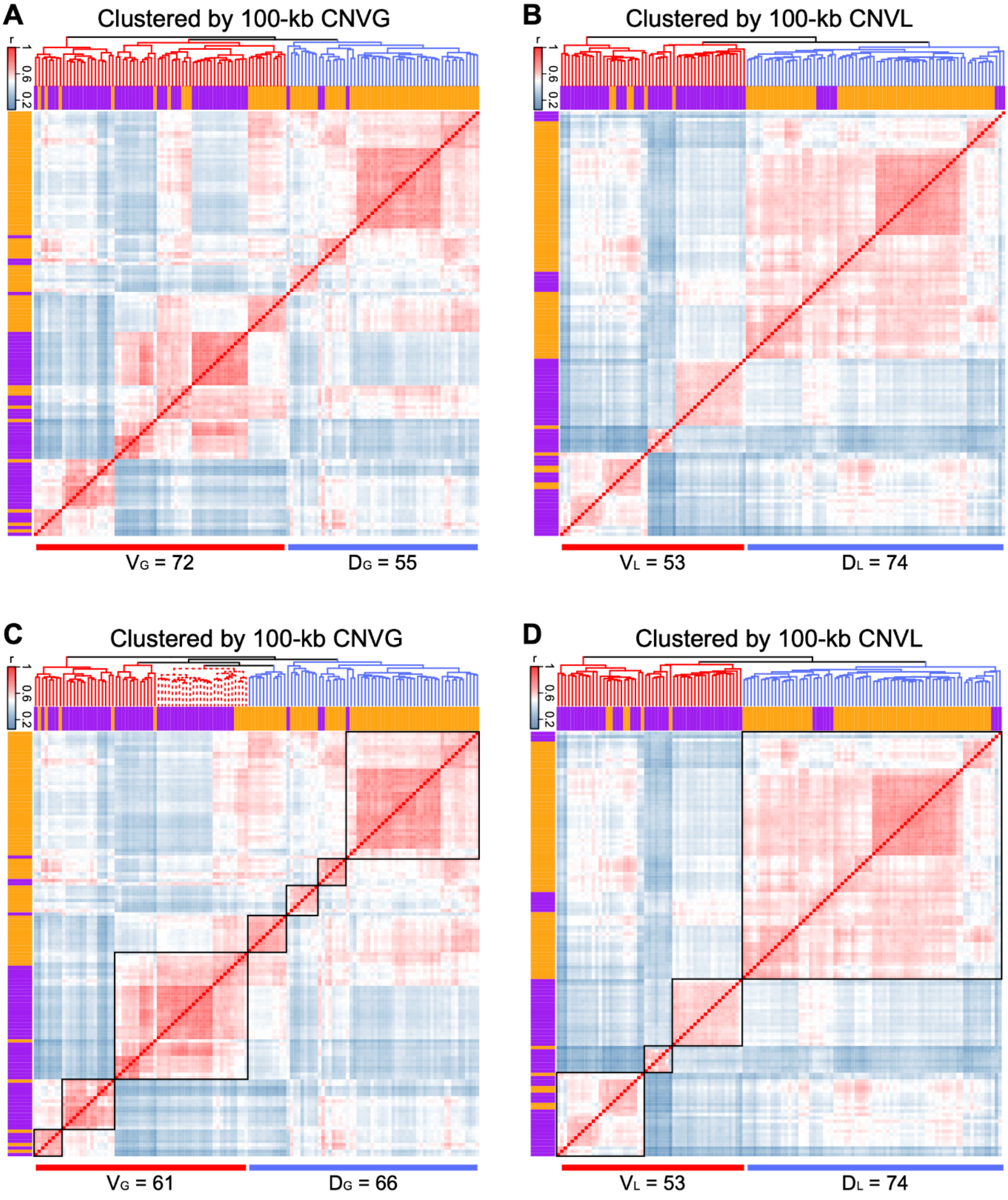


**Figure S2.** Clustering of CNV profiles of 127 P-group samples called from 100-kb sequence windows. The V- and D-subtypes of the 127 samples were obtained based on (A) CNVG profiles using cutree method, (B) CNVL profiles using cutree method, (C) CNVG profiles using semi-supervised method, and (D) CNVL profiles using semi-supervised method. The color of each square in the heat map indicates the correlation coefficient (r) of a pair of samples according to the blue-red thermal scale. The dendrograms on top of the heat maps are bootstrapped for 1,000 times. Branches on the dendrogram were coloured in red for V-type and blue for D-type samples. In the semi-supervised method, the dashed branches on the dendrograms were first rotated around their respective nodes to bring the closely co-localized samples into tightly knit sub-clusters enclosed by black square boxes on the diagonal of each heatmap. Thereupon, all the samples within the same box were all designated as D-type or V-type genomic samples depending on whether the majority clinical subtype of the samples in the box were depression-type or invasion-type.

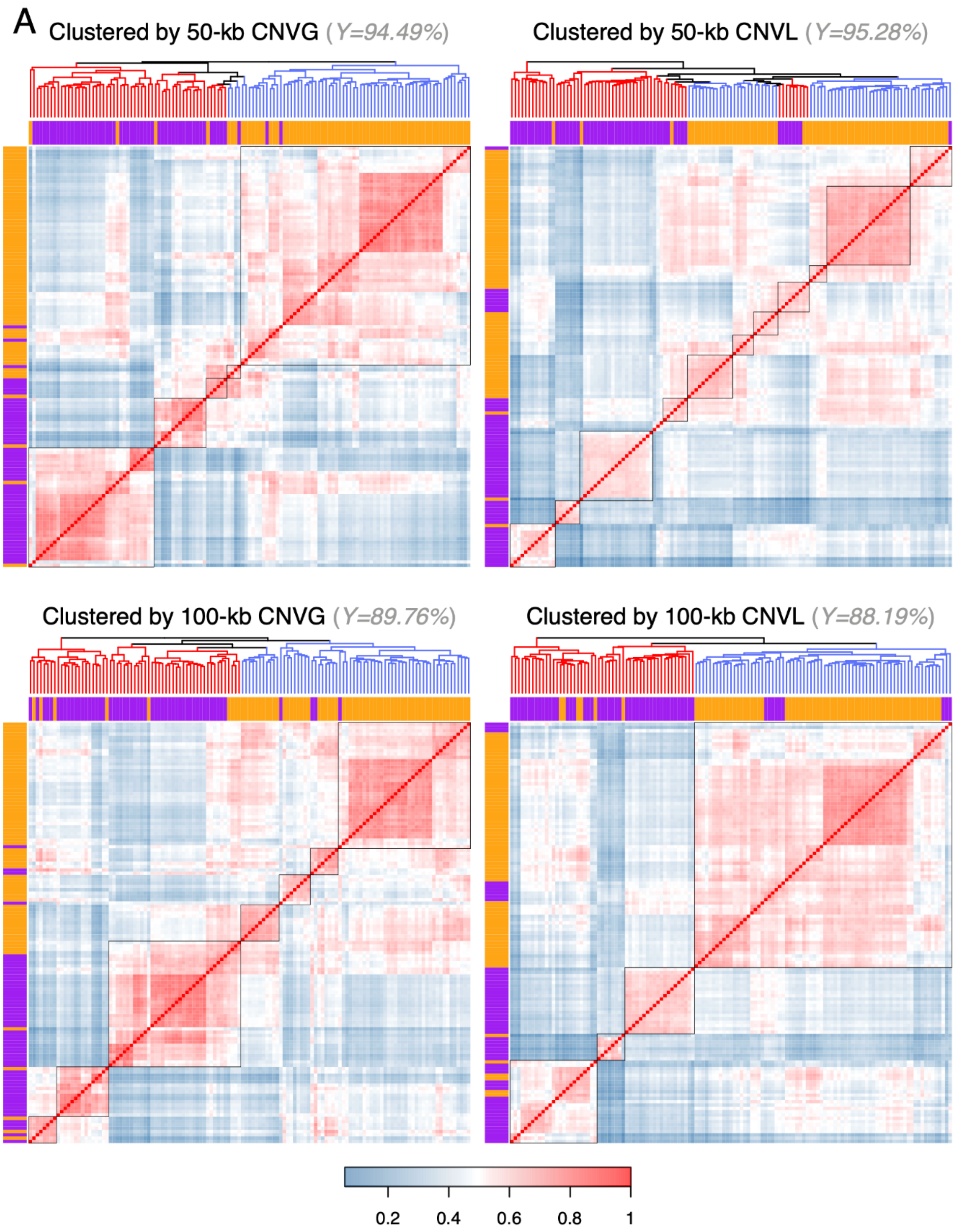


**Figure S3.** (continued on next page)

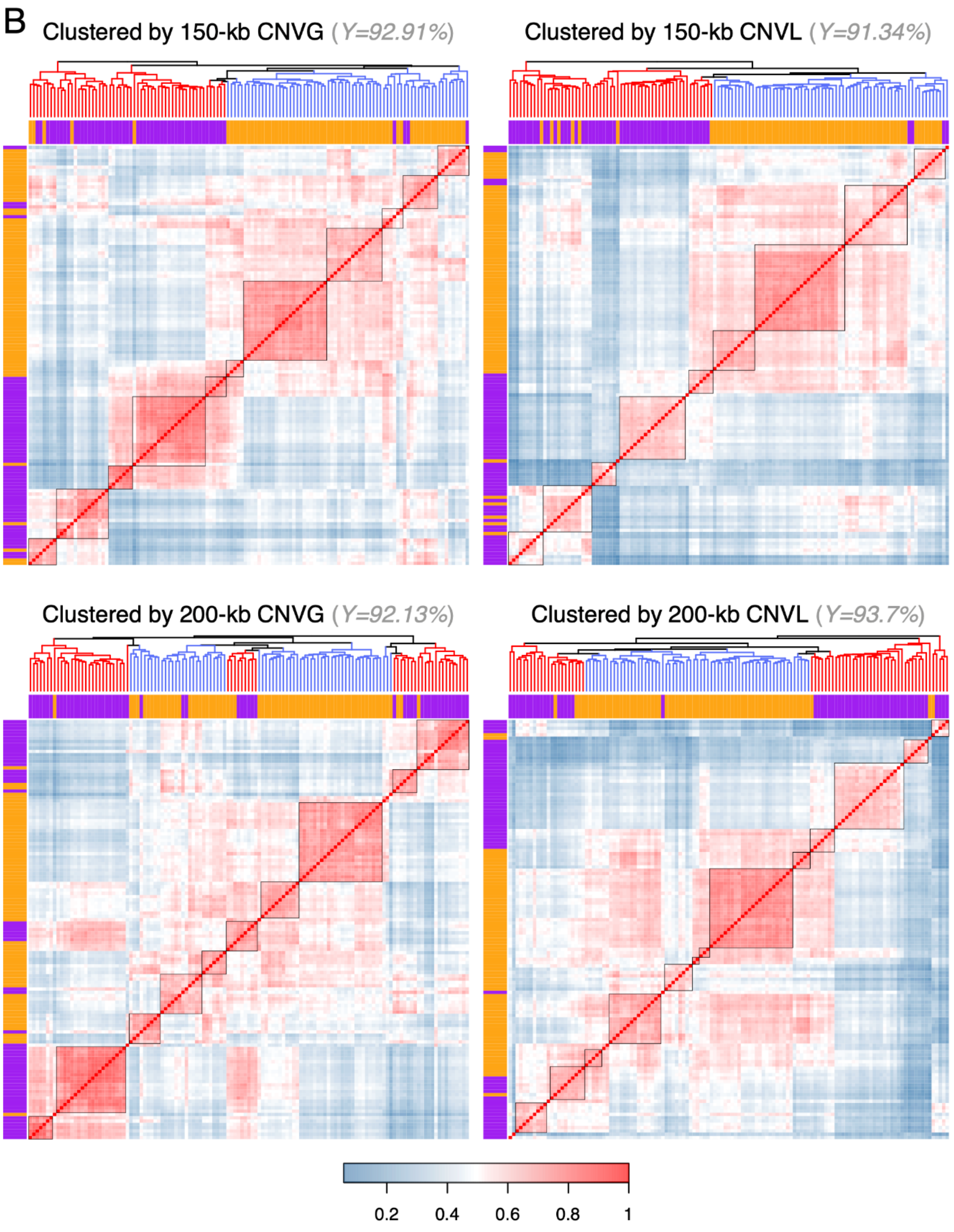


**Figure S3.** (continued on next page)

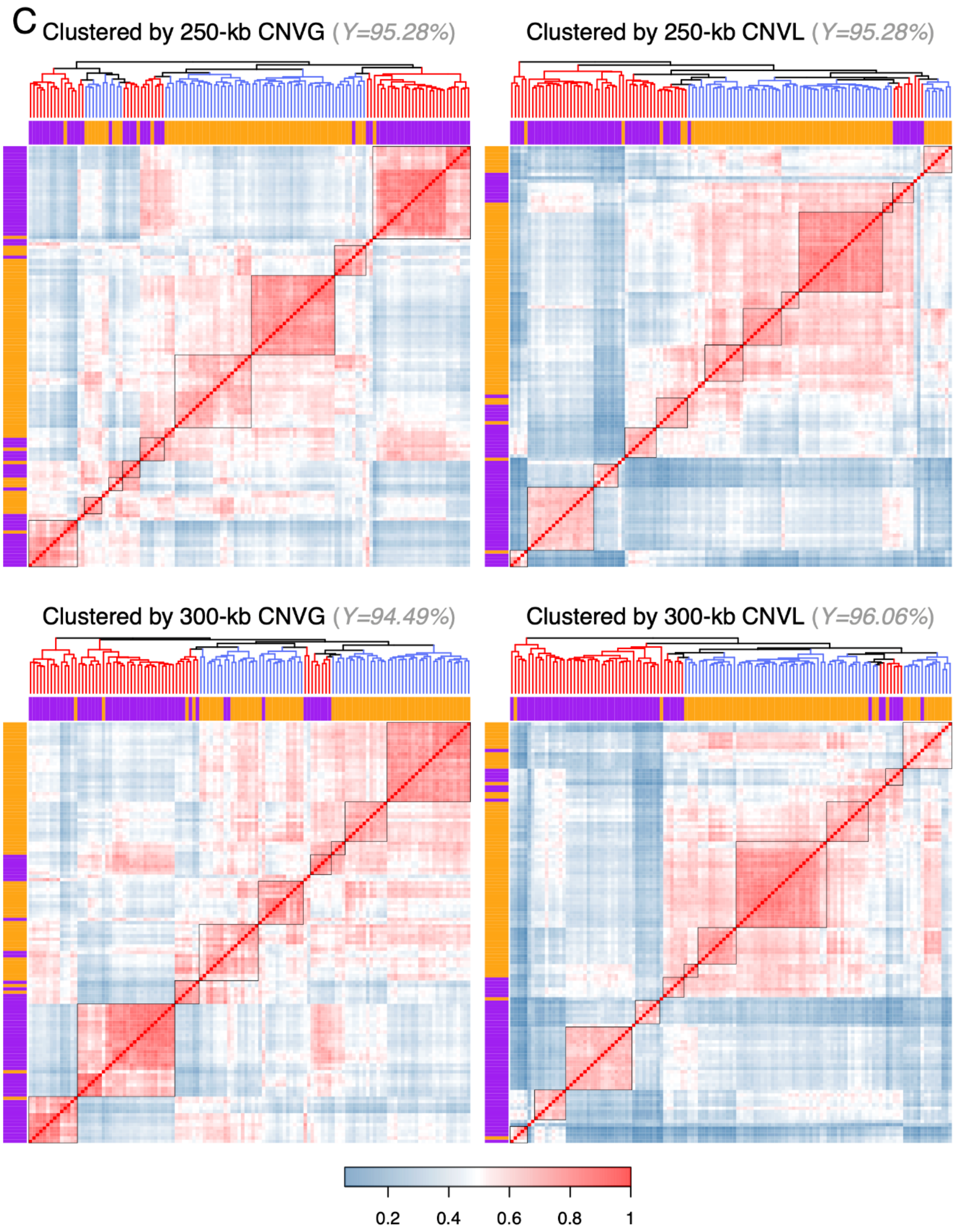


**Figure S3.** (continued on next page)

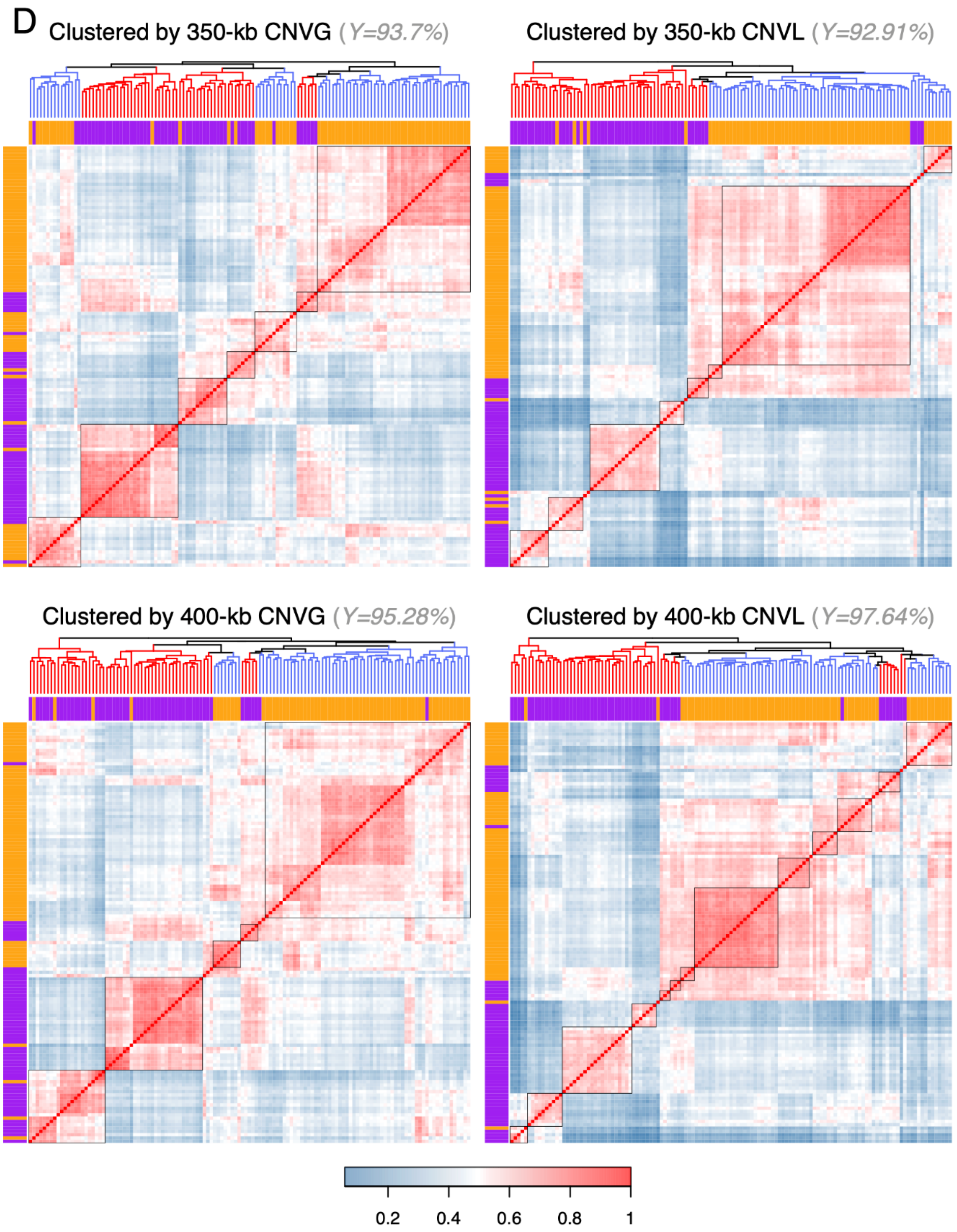


61 **Figure S3.** (continued on next page)

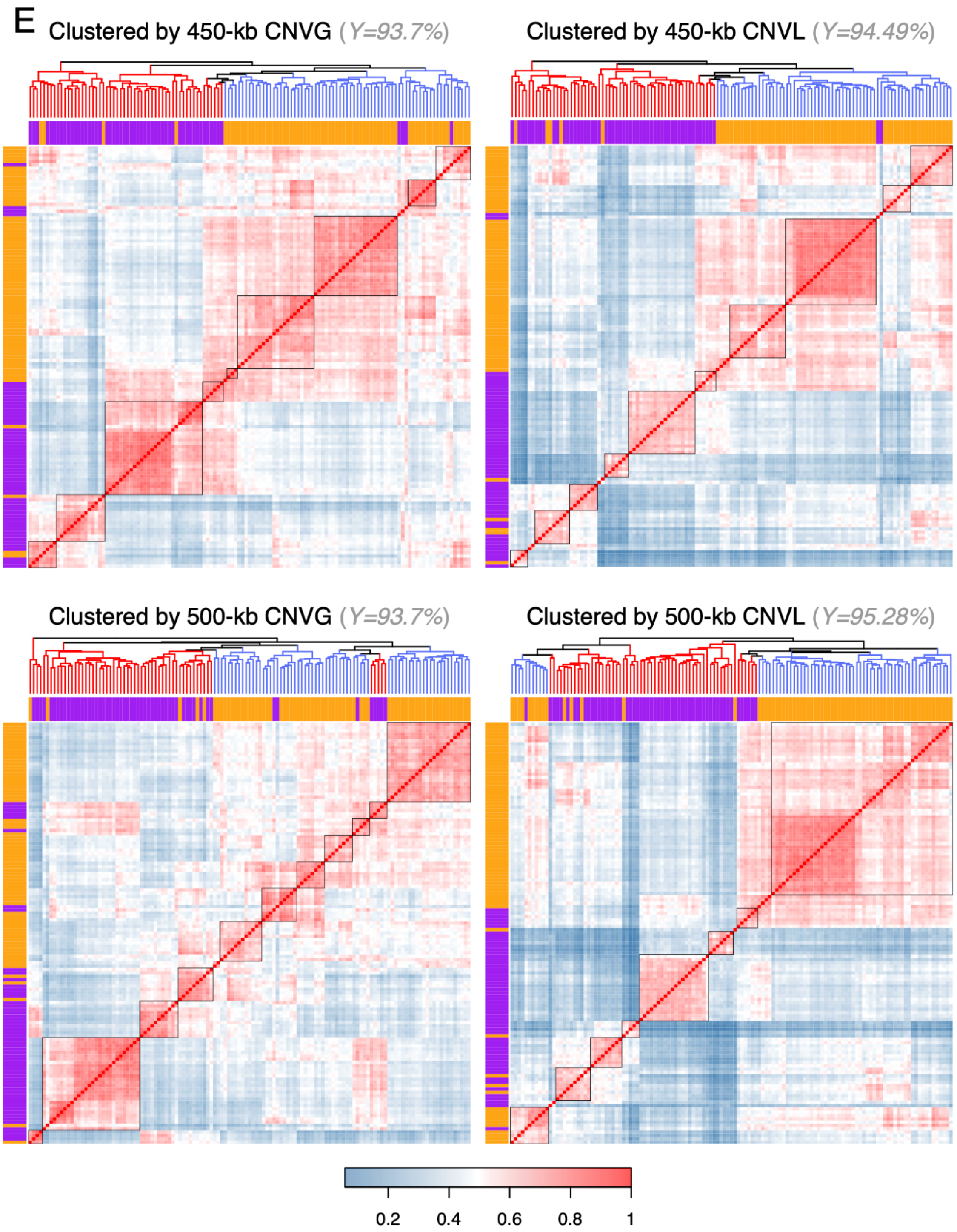


**Figure S3.** Heat maps of correlation coefficients of CNVGs and CNVLs in 127 P-group samples called from (A) 50-kb and 100-kb, (B) 150-kb and 200-kb, (C) 250-kb and 300-kb, (D) 350-kb and 400-kb, (E) 450-kb and 500-kb sequence windows. The color of each square in the heat map indicates the correlation coefficient (r) between a pair of samples shown on the x- and y-axes according to the blue-red thermal scale. The bands at the top of the heat maps represent the subtyping of PMDD samples based on clinical symptoms, with purple bands representing the invasion subtype (n = 56) and orange bands the depression subtype (n = 71). The square black boxes on the diagonal of each heat map, defined groups of samples that showed close correlations with each other in the group, and were therefore uniformly designated as V- or D-type genomes, depending on the majority of the samples displayed an invasion-subtype or depression-subtype of clinical PMDD (see ‘Clustering and grouping of patient samples based on CNV profiles’ in Methods).

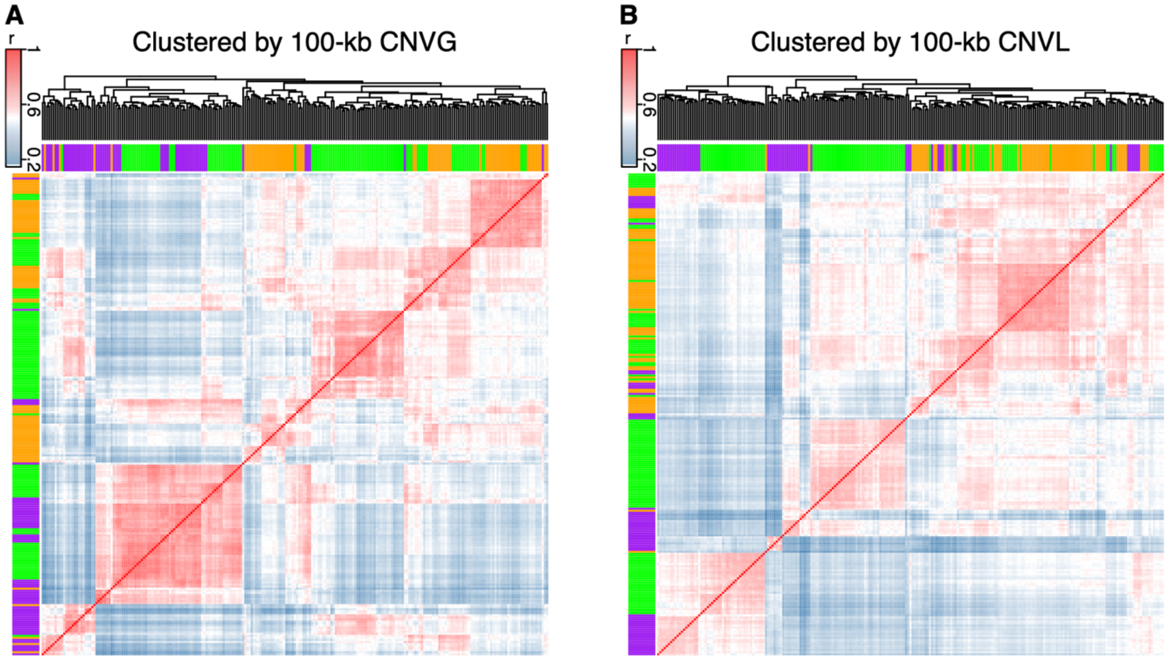


**Figure S4. Hierarchical clustering of CNVGs and CNVLs in a mixture of 127 P-group and 108 C-type samples called from 100-kb sequence windows.** The dendrograms on top of the heat maps were bootstrapped 1,000 times. The color of each square in the heat map indicated the correlation coefficient (r) of a pair of samples according to the blue-red thermal scale. The bands below the dendrograms and on the left-hand side of the heat maps portrayed the subtyping of samples based on clinical symptoms, with purple bands representing the clinically determined invasion subtype (n = 56), orange bands the depression subtype (n = 71), and green bands the control samples (n = 108).

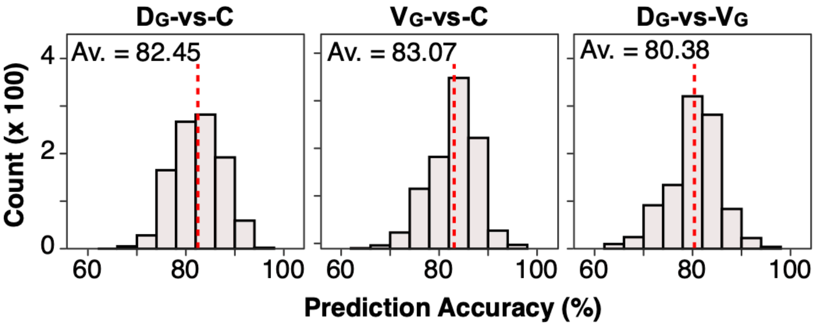


**Figure S5.** The prediction accuracies (estimated using Eqn.2 in Methods) of sample classification using the cutree method in $D_G$-vs-C, $V_G$-vs-C and $D_G$-vs-$V_G$ sample-pairs based on CNV features selected using the correlation method. Subscript G denotes that the D- or V-type samples were derived from the dendrogram of CNVGs (Figure S2A) using the cutree method. For each of the three pairs, prediction accuracy was estimated 1,000 times and the average accuracy (Av.) was given in the pertinent panel. Because the cutree and semi-supervised methods yielded the same sets of classified $D_L$- and $V_L$-type samples, the prediction accuracies obtained with the cutree method were the same as those shown in Figure 2B for the semi-supervised method.

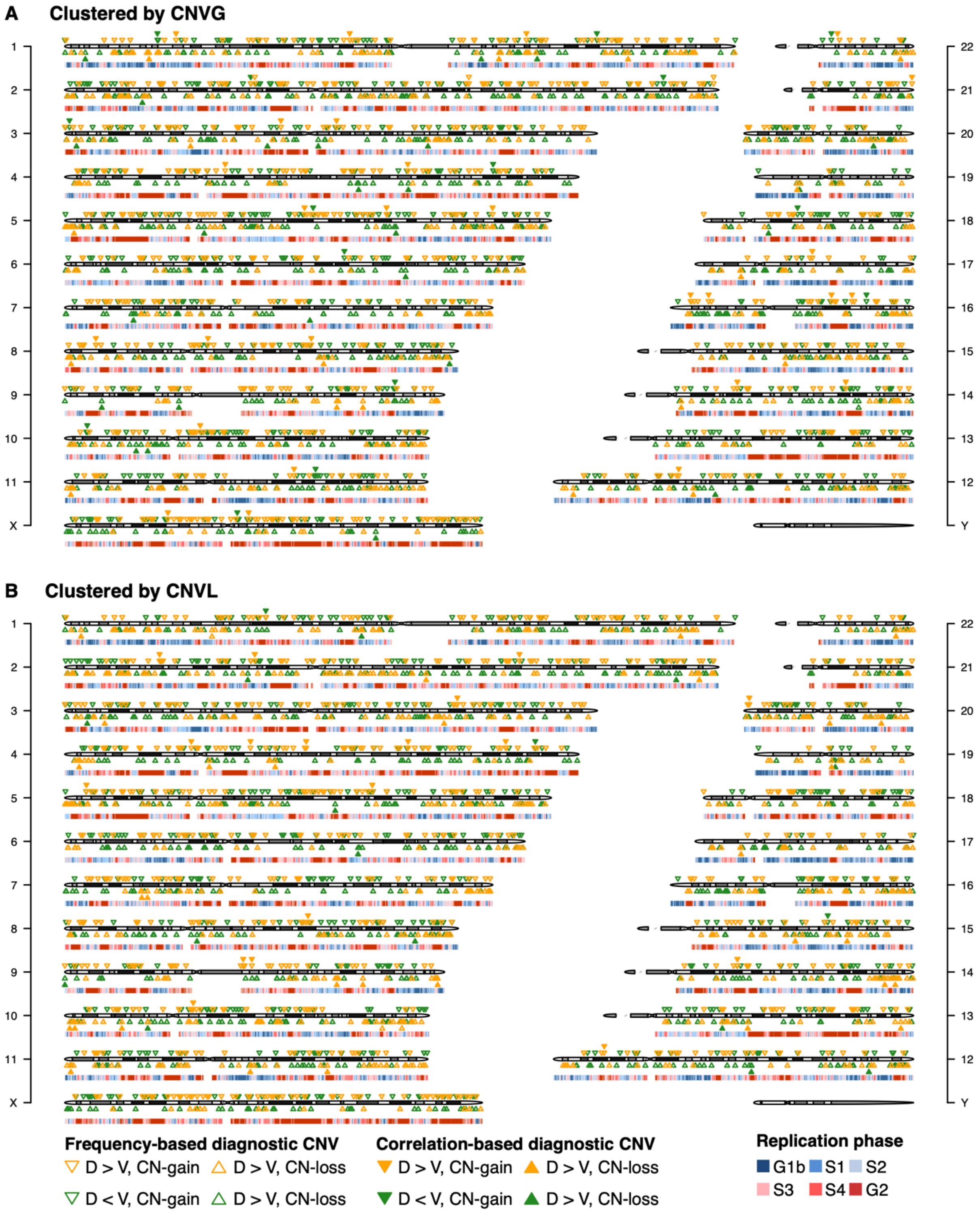


**Figure S6.** Landscape of diagnostic CNV features identified for the D-vs-V comparison. The D- and V-type samples were determined based on the boxed clusters on 100-kb CNV dendrogram delineated by means of the semi-supervised method. The diagnostic CNV features selected for (A) $D_G$-vs-$V_G$ pair and (B) $D_L$-vs-$V_L$ pair are indicated by down-warded triangles for CNVGs and up-warded triangles for CNVLs. Open triangles represent diagnostic CNV features selected using the frequency-based method, and solid triangles represent diagnostic CNV features selected using the correlation-based method. D > V and D < V indicate higher or lower occurrence frequency of D relative to V respectively. The six phases of DNA replication-timing are represented by the colored bands with the early phases (G1b, S1 and S2) in blue, and the late phases (S3, S4 and G2) in red.

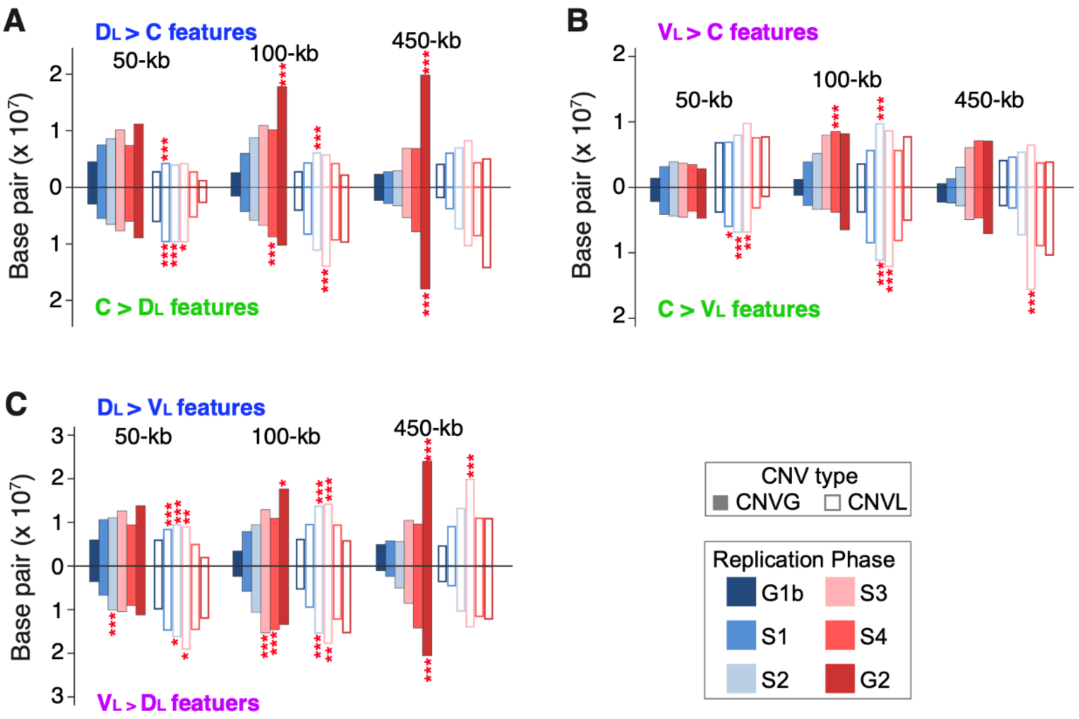


**Figure S7. Distribution of frequency-based diagnostic CNV features among different DNA replication phases.** Number of base pairs of the CNV features called using 50, 100 and 450-kb windows for (A) $D_L$-vs-C, (B) $V_L$-vs-C, and (C) $D_L$-vs-$V_L$ groups. The solid bars represent CNVG features and hollow bars represent CNVL features in each panel. The replication phases G1b to G2 are color coded as shown. The '>' or '<' sign portrays larger or smaller frequencies of the CNV features in favor of the first-named group over the second-named one. L/G represents the ratio of the number of CNVLs over the number of CNVGs. Significant enrichment of CNV features in a particular replication phase in the genome is indicated by asterisks that are color coded according to the replication phase, or in black asterisks for comparison between an L/G value in the upper half of a panel and an L/G value in the lower half (Bonferroni-corrected, *** $p < 0.005$, ** $p < 0.01$, * $p < 0.05$). Numerical $p$-values are shown in Table S14. Subscript L denotes that the D- or V-type samples were derived from the CNVL dendrogram in Figure 1B.

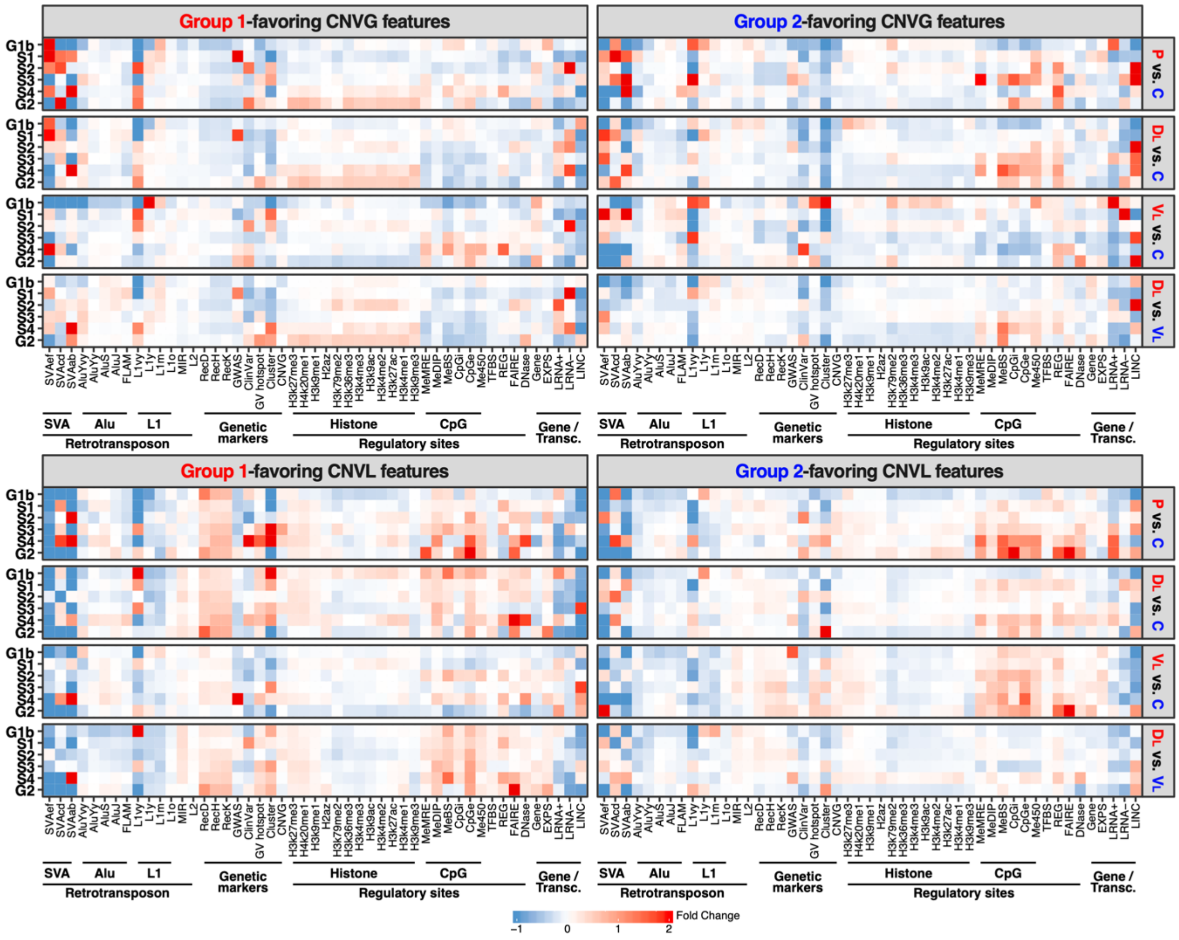


**Figure S8. Enrichment analysis of genomic-feature contents in different replication phases for control and PMDD subtypes**. Frequency-based CNV features diagnostic for C group, i.e., control, as well as that for D and C groups of PMDD samples clustered by CNVL dendrogram, identified with 100-kb scanning windows, were used in the analysis. Enrichment analysis results were plotted for CNVG features in the upper two panels and that for CNVL features in the bottom two panels. Fold-change of each genomic feature in the diagnostic CNV features relative to the non-diagnostic-CNV regions was estimated according to 'Genomic-feature content of diagnostic CNV features in different replication phases' in Methods, and was color-coded based on the thermal scale. Fold-change greater than 2-fold was capped at 2 in the heat map. 'Group 1' indicated the first-named group and 'Group 2' the second-named group in a given pair of samples. Genomic features were grouped into Retrotransposon, Genetic markers, Regulatory sites and Gene/Transcription groups on the x-axis based on their sequence and functional properties. The descriptions of genomic features and numeric data were available in Table S11.

#### Captions of Supplementary Tables

**Table S1.** Clinical diagnosis of all samples analyzed in the present study.

**Table S2.1** CNVG profiles of all samples with CNV sizes ranging from 50kb to 500kb.

**Table S2.2** CNVL profiles of all samples with CNV sizes ranging from 50kb to 500kb.

**Table S3.1** CNV-based typing of PMDD genomes using cutree method.

**Table S3.2** CNV-based typing of PMDD genomes using semi-supervised method.

**Table S4.1** Consistency between clinical subtyping of PMDD cases and CNV-based typing of genomes using cutree method.

**Table S4.2** Consistency between clinical subtyping of PMDD cases and CNV -based typing of genomes using semi-supervised method.

**Table S5.1** Functional annotation of genes that overlapped with frequency-based diagnostic CNV features for distinguishing between the D- and V-types using cutree method.

**Table S5.2** Functional annotation of genes that overlapped with frequency-based diagnostic CNV features for distinguishing between the D- and V-types using semi-supervised method.

**Table S6.** Number of windows randomly selected in Monte Carlo simulations.

**Table S7.1** Information on frequency-based and correlation-based diagnostic CNV features for distinguishing the D- and V-types using cutree method.

**Table S7.2** Information on frequency-based and correlation-based diagnostic CNV features for distinguishing D- and V-type classifications semi-supervised based on clinical diagnosis.

**Table S8.** List of correlation-based CNV features useful for distinguishing between D and V subtypes.

**Table S9.** Shared CNV features among P-vs-C, D-vs-C and V-vs-C comparisons.

**Table S10.** KEGG pathways enriched in 50-, 100- and 450-kb frequency-based CNV features with < 0.05 empirical $p$-values.

**Table S11.** Descriptions and distributions of genomic features in 100-kb frequency-based CNV features in different replication phases.

**Table S12.** Number of frequency-based CNV features jointly identified by the cutree and semi-supervised methods.

**Table S13.** Comparison of replication phase-distributions of frequency-based CNV features identified using the cutree and semi-supervised methods.

**Table S14.** Statistics for enrichment of CNV features in different replication phases.

**Table S15.** Genetic distance estimation for different pairs of C, P, D and V types.
